# Renal proximal tubules are sensitive to metabolic acidosis

**DOI:** 10.1101/2024.08.19.608649

**Authors:** J. Christopher Hennings, Keerthana S. Murthy, Nicolas Picard, Inês Cabrita, David Böhm, Maria E. Krause, Vandit Shah, Jennifer Baraka-Vidot, Mukhran Khundadze, Tobias Stauber, Detlef Böckenhauer, Thomas J. Jentsch, Sebastian Bachmann, Bernhard Schermer, Dominique Eladari, Régine Chambrey, Christian A. Hübner

## Abstract

Patients suffering from distal renal tubular acidosis (dRTA) are sometimes diagnosed with proximal tubule dysfunction with leaks of phosphate, uric acid, amino acids, and low-molecular-weight proteins, also known as Fanconi-like syndrome. The underlying molecular basis is largely elusive. We previously reported on *Atp6v0a4* knockout (KO) mice, which exhibit severe metabolic acidosis in combination with proximal tubule dysfunction as evidenced by phosphaturia and proteinuria. Here, we show that Rab7, a key regulator of endo-lysosomal trafficking and lysosomal biogenesis, is strongly diminished in proximal tubules of *Atp6v0a4* KO mice, while the number of abnormal Ist1-labelled Lamp1-positive vesicles is increased. This is accompanied by the accumulation of autophagosomes, autolysosomes and autophagic substrates. Importantly, correction of metabolic acidosis with bicarbonate therapy resolves proximal tubule dysfunction and trafficking defects in *Atp6v0a4* KO mice. Acid-challenged wildtype mice also show trafficking defects with Rab7-downregulation and an increase in Ist1-labeled Lamp1-positive vesicles and develop proximal tubule damage in the long-term. Similar acidosis-induced alterations also occur in human kidney organoids. Altogether, our data provide insights, why patients suffering from severe dRTA may develop a Fanconi-like syndrome, which may contribute to the progression of chronic kidney failure.

**Translational Statement:** Patients with renal acidosis caused by impaired proton secretion in the collecting duct (distal renal tubular acidosis - dRTA) sometimes show unexplained symptoms of proximal tubule dysfunction such as proteinuria and phosphaturia. Here, we show that proximal tubules are particularly sensitive to acidosis as evidenced by impaired trafficking, lysosomal damage and accumulation of autophagic substrates. We also show that early treatment of dRTA by alkali supplementation can prevent proximal tubule dysfunction. Because metabolic acidosis represents a well-known risk factor for the progression of chronic kidney disease (CKD), our findings highlight the potential clinical importance of early alkali supplementation to delay disease progression.

## Introduction

Most molecules in our body show a remarkable sensitivity to changes in pH. Therefore, a tight control of acid-base homeostasis and pH is critical. In our organism, HCO_3_^−^/CO_2_ is the most important buffer, because CO_2_ can be regulated by respiration and HCO_3_^−^ by the kidneys. The kidneys reabsorb virtually all filtered HCO_3_^−^ and additionally generate new HCO_3_^−^ by net H^+^ excretion. The largest fraction is reclaimed in the proximal tubule by H^+^ excretion via the apical Na^+^-H^+^-exchanger. If this process is disturbed, the resulting HCO_3_^−^ loss causes proximal or type II renal tubular acidosis (pRTA). In the distal nephron, renal bicarbonate regeneration and H^+^ excretion are mediated by intercalated cells. Here, a defect of distal H^+^ excretion results in distal or type I renal tubular acidosis (dRTA), which can be either caused by mutations of the V-ATPase subunits ATP6V0A4 or ATP6V1B1, the basolateral anion-exchanger SLC4A1 (AE1), the forkhead transcription factor FOXI1 ^1^ or the β-propeller containing protein WDR72 ^2^. Remarkably, patients with either pRTA or dRTA can suffer from proximal tubule dysfunction with leaks of phosphate, uric acid, amino acids, and low-molecular-weight proteins, also known as Fanconi-like syndrome ^3–6^.

We previously reported on a mouse model for ATP6V0A4-related dRTA ^7^. Our *Atp6v0a4* knockout (KO) mice suffer from severe metabolic acidosis and proximal tubule dysfunction as evidenced by phosphaturia and proteinuria and die within the first weeks of life. Because the Atp6v0a4 subunit is not only expressed in intercalated cells but also in proximal tubule cells ^8^, proximal tubule dysfunction upon disruption of *Atp6v0a4* may be cell intrinsic. Remarkably, however, another independent study did not report proximal tubule dysfunction in adult *Atp6v0a4* KO mice, which had been rescued by chronic HCO_3_^−^ supplementation ^9^.

Here, we provide clinical data from dRTA patients, who presented with a Fanconi-like syndrome during acidotic episodes. A Fanconi-like syndrome was also observed in acidotic *Atp6v0a4* KO mice and resolved upon bicarbonate supplementation. Proximal tubule cells of acidotic *Atp6v0a4* KO mice displayed severe trafficking defects with alterations of the degradative pathway. Such alterations were absent in mosaic proximal tubule cell-specific KO mice, which did not display acidosis. Similar trafficking defects also developed in proximal tubules of acid-challenged wildtype (WT) mice and human kidney organoids. Taken together, our models provide insights into the underlying pathophysiology of acidosis-induced Fanconi-like syndrome.

## Methods

### Human patients

Clinical data was extracted from previous studies ^3, 10^.

### Mice

Constitutive *Atp6v0a4* KO mice were described in ^7^. Conditional *Atp6v0a4* KO were generated by crossing mice with the targeted *Atp6v0a4* locus ^7^ to Flpe-Deleter mice (**Supplementary Figure S8**). Pepck-cre mice were described in ^11^. ApoE-cre mice were described in ^12^. All mouse strains were backcrossed with C57BL/6 mice for at least 10 generations. Male C57BL/6 mice were obtained from Janvier and habituated for 2 weeks before experiments. The experimental unit of our study is the single mouse. Except for animal welfare reasons, no mice were excluded from our study.

### Bicarbonate rescue of *Atp6v0a4* KO mice

Pregnant *Atp6v0a4*^+/-^ mice were randomly selected for treatment with 0.14 M sodium bicarbonate (Carl Roth) or with regular tap water. Litters with male and female pups were genotyped and *Atp6v0a4*^+/+^ and *Atp6v0a4*^-/-^ mice were euthanized at 3 weeks of age (**Figure 1a**). Because of variations in litter size and increased mortality of untreated Atp6v0a4^-/-^ mice, the sample size for each cohort were not always identical as indicated. Overall, we analyzed n = 23 water-treated *Atp6v0a4*^+/+^, n = 19 *Atp6v0a4*^-/-^ and n = 19 bicarbonate-treated *Atp6v0a4*^+/+^, n = 19 *Atp6v0a4*^-/-^ mice. For physiological measurements, we aimed for a minimum of 5 mice per cohort, for protein isolation 6 mice per cohort and for immunofluorescence analysis 3 mice per cohort. Sample sizes depended on animal availability, our previous experience with comparable experiments and legal limitations because of the granted animal licenses.

**Figure 1:**
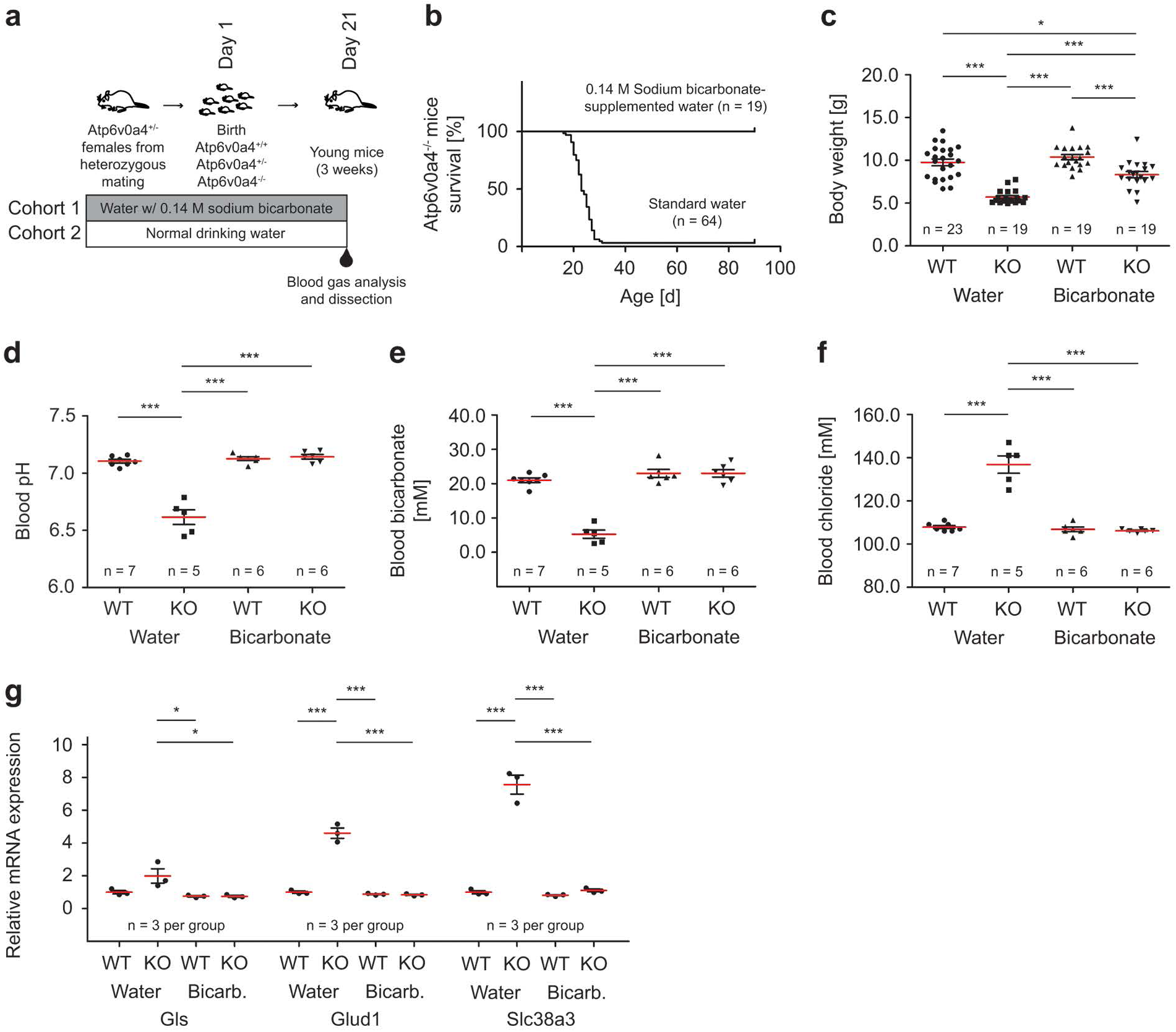
Severe metabolic acidosis in *Atp6v0a4* KO mice is rescued by bicarbonate supplementation. **(a)** Schematic representation of the bicarbonate rescue experiment. **(b)** Kaplan-Meier plot for survival of *Atp6v0a4* KO mice with and without bicarbonate supplementation. Without bicarbonate supplementation, more than 98% of *Atp6v0a4* KO died before weaning. n = number of mice. (c) Bicarbonate supplementation partially rescues body weight loss in *Atp6v0a4* KO mice. n = number of mice. Each data point represents one mouse. One-way ANOVA with Tukey·s multiple comparison test. **(d-f)** *Atp6v0a4* KO mice display severe metabolic acidosis as shown by blood pH, blood bicarbonate and blood chloride at 21 days of age, which is rescued by bicarbonate supplementation. n = number of mice. Each data point represents one mouse. One-way ANOVA with Tukey·s multiple comparison test. **(g)** Quantitative RT-PCR of whole kidney mRNA from 21-day-old mice shows increased expression of genes involved in ammoniagenesis: Glutaminase (Gls), Glutamate dehydrogenase 1 (Glud1), Solute carrier family 38 member 3 (Slc38a3). n = 3 mice per cohort. Each data point represents one mouse. One-way ANOVA with Tukey·s multiple comparison test. Quantitative data are represented as mean± s.e.m. with individual data points. *\*P* < 0.05; *\*\*P* < 0.005; *\*\*\*P* < 0.0005.

### Acid-loading of C57BL/6 mice

For acid-loading of C57BL/6 mice, mice had free access to tap water supplemented with 0.28 M NH_4_Cl (Carl Roth). Mice were euthanized for organ removal at 3 months of age. For this experiment, we aimed at 10 mice per cohort. Because some animals died unexpectedly during the final blood sampling the experiment was finished with less mice. For physiological measurements, we aimed for a minimum of 5 mice per cohort, for RNA/protein isolation 4 mice per cohort and for immunofluorescence analysis 3 mice per cohort.

### Statistics

Data are presented as mean ± s.e.m. unless otherwise indicated. The sample number (n) indicates the number of mice or organoids. Single data points indicate the number of analyzed proximal tubules in immunofluorescence experiments. We used two-sided Student’s t-test or one-way analysis of variance (ANOVA) followed by Tukeýs multiple-comparison tests after testing for normal distribution. Prism v.5.0 was used to generate graphs and calculate statistics.

### Other methods

All other methods are found in the Supplementary Methods.

## Results

### Severe dRTA is associated with proximal tubule dysfunction in humans

Transient proximal tubular dysfunction has been repeatedly reported in patients with dRTA ^3, 6, 13–15^. Here, we provide examples from Great Ormond Street Hospital for Children in London, which have been partially published previously (**Table 1**). Importantly, the proximal dysfunction largely resolved with correction of acidosis.

**Table 1:** Biochemical details of patients with dRTA and transient proximal tubular dysfunction. Acidosis manifested within the first year of life and is reflected by decreased plasma bicarbonate concentrations. In patient 4, acidosis re-occurred at 14 years of age, when the medication was discontinued. During acidosis, patients 1-4 showed hypophosphatemia with renal phosphate wasting and low molecular weight proteinuria (LMWP). Patient 5 exhibited LMWP only, reflecting the wide spectrum of the severity of proximal tubule dysfunction. Correction of the acidosis normalized biochemical parameters, although patients 1 and 3 had persistent borderline LMWP, potentially indicating a permanent proximal tubule defect. All patients were seen at Great Ormond Street Hospital for Children in London and clinical and/or genetic details of some of these patients have been published previously. All reported variants were homozygous, except for patient 2. Parameters shown: plasma bicarbonate concentration; plasma phosphate concentration; TmP/GFR: tubular maximum reabsorption of phosphate corrected for glomerular filtration rate; urine Retinol-Binding Protein (RBP) corrected for creatinine. Increased values in gray.

| Patient | 1 |  | 2 |  | 3 |  | 4 |  | 5 |  |
| --- | --- | --- | --- | --- | --- | --- | --- | --- | --- | --- |
| Gene | *ATP6V1B1* |  | *ATP6V1B1* |  | *ATP6V1B1* |  | *ATP6V0A4* |  | *ATP6V0A4* |  |
| Variant | c.905G>C<br>c.1469C>T<br>p.(R320P)<br>p.(P490L) |  | c.1181G>A<br>p.(R394Q) |  | c.1312_1315del<br>p.(D483Mfs*13) |  | c.1107delC<br>p.(N370Ifs*22) |  | c.2257+1G>A<br>p.(?) |  |
| Acidosis | + | - | + | - | + | - | + | - | + | - |
| Plasma HCO<sub>3</sub><sup>-</sup><br>(normal >20)<br>[mmol/l] | 14 | 21 | 12 | 26 | 12 | 24 | 13 | 27 | 12 | 23 |
| Plasma PO<sub>4</sub><sup>3-</sup><br>(normal >1.2)<br>[mmol/l] | 0.96 | 1.26 | 1.05 | 1.64 | 0.5 | 1.32 | 0.92 | 2.28 | 1.87 | 2.17 |
| TmP/GFR<br>(normal >1.15)<br>[mmol/l] | 0.56 | 1.55 | 0.35 | 1.45 | 0.35 | 1.2 | 0.65 | n/a | n/a | n/a |
| Urine RBP/creatinine<br>(normal <100)<br>[µg/mmol] | 34000 | 126 | 17750 | 25 | 10900 | 164 | 8975 | 87 | 22300 | 86 |

### *Atp6v0a4* KO mice suffer from severe metabolic acidosis, which is largely alleviated by bicarbonate supplementation

We mated heterozygous *Atp6v0a4*^+/-^ mice to obtain WT (*Atp6v0a4*^+/+^) and KO (*Atp6v0a4*^-/-^) littermates. Pregnant mice received either regular tap water or tap water supplemented with 0.14 M sodium bicarbonate, which was continued after delivery (**Figure 1a**). While more than 98% of untreated *Atp6v0a4* KO mice did not survive beyond 30 days after birth, bicarbonate-supplementation rescued KO mice (**Figure 1b**). Bicarbonate supplementation also improved the postnatal weight gain of *Atp6v0a4* KO mice (**Figure 1c**). While blood gas analyses at postnatal day 21 confirmed severe hyperchloremic metabolic acidosis in untreated *Atp6v0a4* KO mice, the acid-base status was normalized in *Atp6v0a4* KO mice supplemented with bicarbonate (**Figure 1d-f**). As evidenced by qPCR of whole kidney RNA, key molecules required for ammoniagenesis were dramatically upregulated in untreated, but not in bicarbonate-supplemented *Atp6v0a4* KO mice confirming that the latter had a normal acid-base status (**Figure 1g**).

These data demonstrate that bicarbonate supplementation is an effective strategy to correct metabolic acidosis caused by *Atp6v0a4* deficiency.

### Bicarbonate supplementation prevents the Fanconi-like syndrome in *Atp6v0a4* KO mice

Like patients with a Fanconi-like syndrome, *Atp6v0a4* KO mice displayed hyperphosphaturia (**Figure 2a**) and proteinuria (**Figure 2b-c**) with albuminuria (**Figure 2d**), which could be prevented by bicarbonate supplementation (**Figure 2a-d**). The hyperphosphaturia can be explained by the dramatically decrease of apical expression of the Na^+^-phosphate co-transporter Slc34a1 (NaPi2a) (**Figure 2e**). Histological (**Supplementary Figure S1a)** and electron microscopy analysis (**Supplementary Figure S1b**) suggested that the typical brush border of proximal tubular cells remained intact. The labeling for Slc34a1 at the brush border was identical to WT in bicarbonate-supplemented *Atp6v0a4* KO mice (**Figure 2e**).

**Figure 2:**
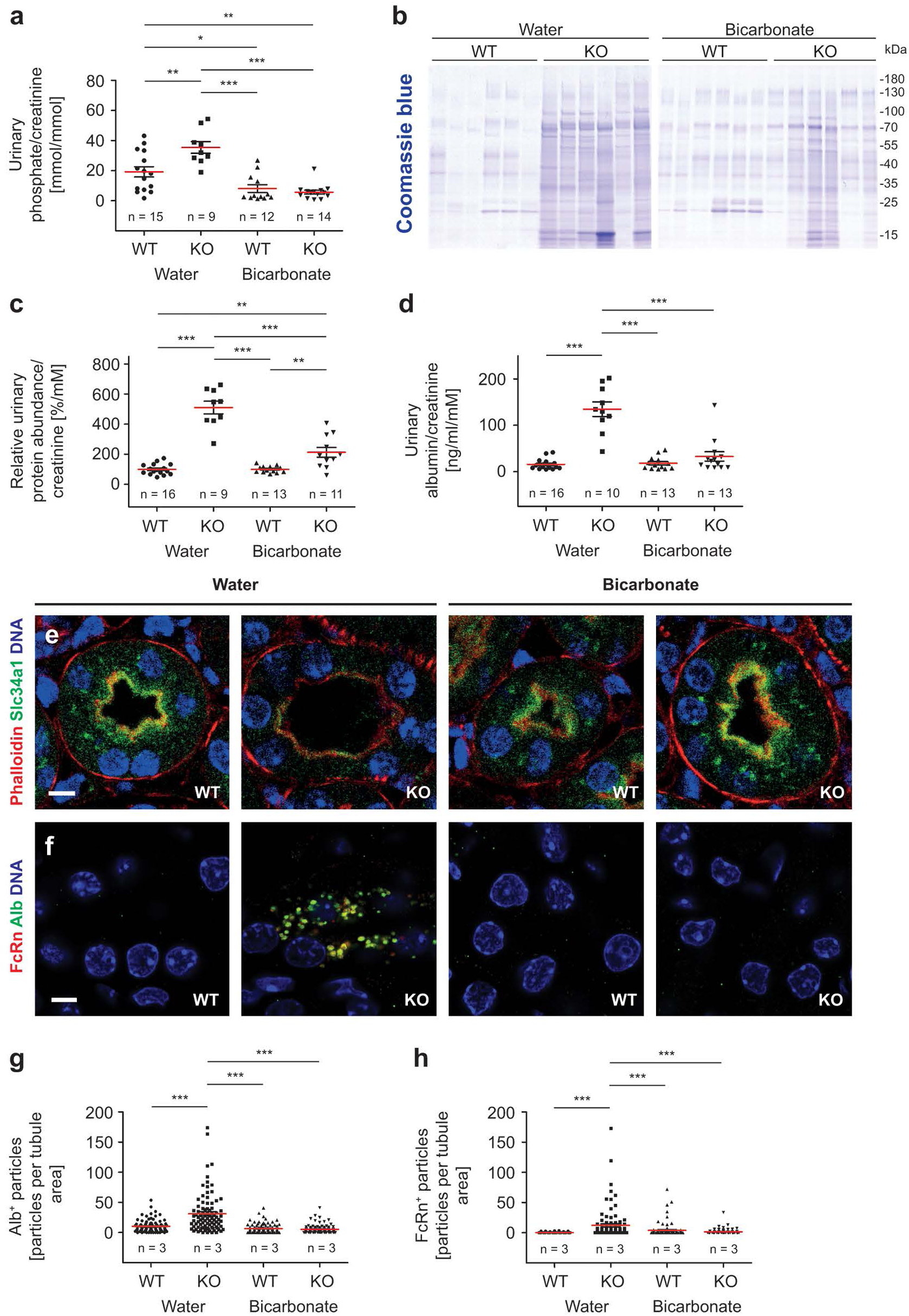
The Fanconi-like syndrome of *Atp6v0a4* KO mice is prevented by bicarbonate supplementation. **(a)** Urinary phosphate excretion normalized to creatinine is increased in 21-day-old *Atp6v0a4* KO mice compared to WT mice. Bicarbonate supplementation reduces phosphate excretion independent of the genotype. n = number of mice. Each data point represents one mouse. One-way ANOVA with Tukey’s multiple comparison test. **(b)** Coomassie-stained Western blot. Urinary protein excretion normalized to creatinine is increased in 21-day-old *Atp6v0a4* KO mice compared to WT mice. Bicarbonate supplementation partly rescues urinary protein excretion in *Atp6v0a4* KO mice. Each lane represents a urine sample from one individual mouse. (c) Quantification of urinary proteins from the Coomassie-stained Western blot. n = number of mice. Each data point represents one mouse. One-way ANOVA with Tukey’s multiple comparison test. **(d)** Urinary albumin excretion normalized to creatinine is increased in untreated 21-day-old *Atp6v0a4* KO mice and normalizes upon bicarbonate supplementation. n = number of mice. Each data point represents one mouse. One-way ANOVA with Tukey’s multiple comparison test. **(e)** The expression of the apical phosphate transporter Slc34a 1 is reduced in 21-day-old *Atp6v0a4* KO mice, but normalizes upon bicarbonate supplementation. Scale bar 5 µm. **(f)** Vesicles positive for Albumin- and FcRn accumulate in proximal tubule cells in the medulla of untreated 21-day-old *Atp6v0a4* KO mice, but not in WT and/or in bicarbonate-treated mice. Single channels are shown in supplemental figures. Scale bar 5 µm. **(g,h)** Quantification of particles positive for Alb and FcRn per proximal tubule area. Both numbers of Albumin- and FcRn-positive particles are increased in untreated *Atp6v0a4* KO mice. Bicarbonate supplementation lowers particles to WT levels. n = number of mice. Each data point represents one proximal tubule. One-way ANOVA with Tukey’s multiple comparison test. Quantitative data are represented as mean ± s.e.m. with individual data points. *\*P* < 0.05; *\*\*P* < 0.005; *\*\*\*P* < 0.0005.

Normally, filtered proteins including albumin are reabsorbed by proximal tubule cells via receptor-mediated endocytosis ^16^. After endocytosis, albumin is either degraded upon fusion of endosomes with lysosomes, or transcytosed upon binding to the neonatal Fc receptor (FcRn) ^17^. When kidney sections from *Atp6v0a4* KO mice were stained for albumin and FcRn, we found an accumulation of both FcRn-positive and -negative vesicles filled with albumin in all segments of proximal tubules in *Atp6v0a4* KO mice (**Figure 2f-h, Supplementary Figure S2a**) supporting the finding that straight segments participate in protein uptake during proteinuric stress ^18^. Accordingly, we observed an accumulation of basophilic most likely protein-loaded vesicles in Toluidine blue-stained kidney sections from *Atp6v0a4* KO mice (**Supplementary Figure S2b**). Such vesicles were absent in WT and bicarbonate supplemented KO mice (**Figure 2f-h**).

We conclude that bicarbonate supplementation prevents the Fanconi-like syndrome observed in *Atp6v0a4* KO mice.

### The endocytic pathway is severely compromised in untreated *Atp6v0a4* KO mice

The accumulation of albumin-positive vesicles in *Atp6v0a4* KO mice prompted us to study the endocytic pathway, which delivers macromolecules for degradation to lysosomes through several endosomal intermediates. Signals for the multiligand endocytic receptor Megalin (Lrp2), which largely mediates the uptake of filtered proteins by proximal tubule cells ^17^, did not differ between genotypes and overlapped with the early endosome marker Eea1 (**Supplementary Figure S1a**). Early endosomes, which also label for the small GTPase Rab5 ^19^, sort material for recycling or degradation. During sorting processes, endocytic intermediates are remodeled into later stage endosomes, which involves several kiss-and-run and fusion events ^20^. The resulting late endosomes can either release their content to the extracellular space or fuse with lysosomes. This maturation process is regulated by a sequential shift of activity from the early endosomal Rab5 to the late endosomal Rab7 ^21, 22^. While the total abundance of Eea1 was not changed in whole kidney protein lysates, it was reduced for Rab5 (**Figure 3a**). Staining kidney sections for Rab5 further revealed that the sharp peak for Rab5 signals below the brush border was flattened in KO samples (**Figure 3b-c, Supplementary Figure S3a-b**). Suggesting that the maturation of late endosomes may be compromised, Rab7 abundance was also strongly diminished in kidney lysates (**Figure 3a**) and Rab7 labeled puncta drastically reduced in proximal tubule sections of untreated *Atp6v0a4* KO mice (**Figure 3d-e**). Both the overall abundance as well as the number of Rab7 labeled puncta did not differ from controls in bicarbonate-supplemented mice (**Figure 3a and e**).

**Figure 3:**
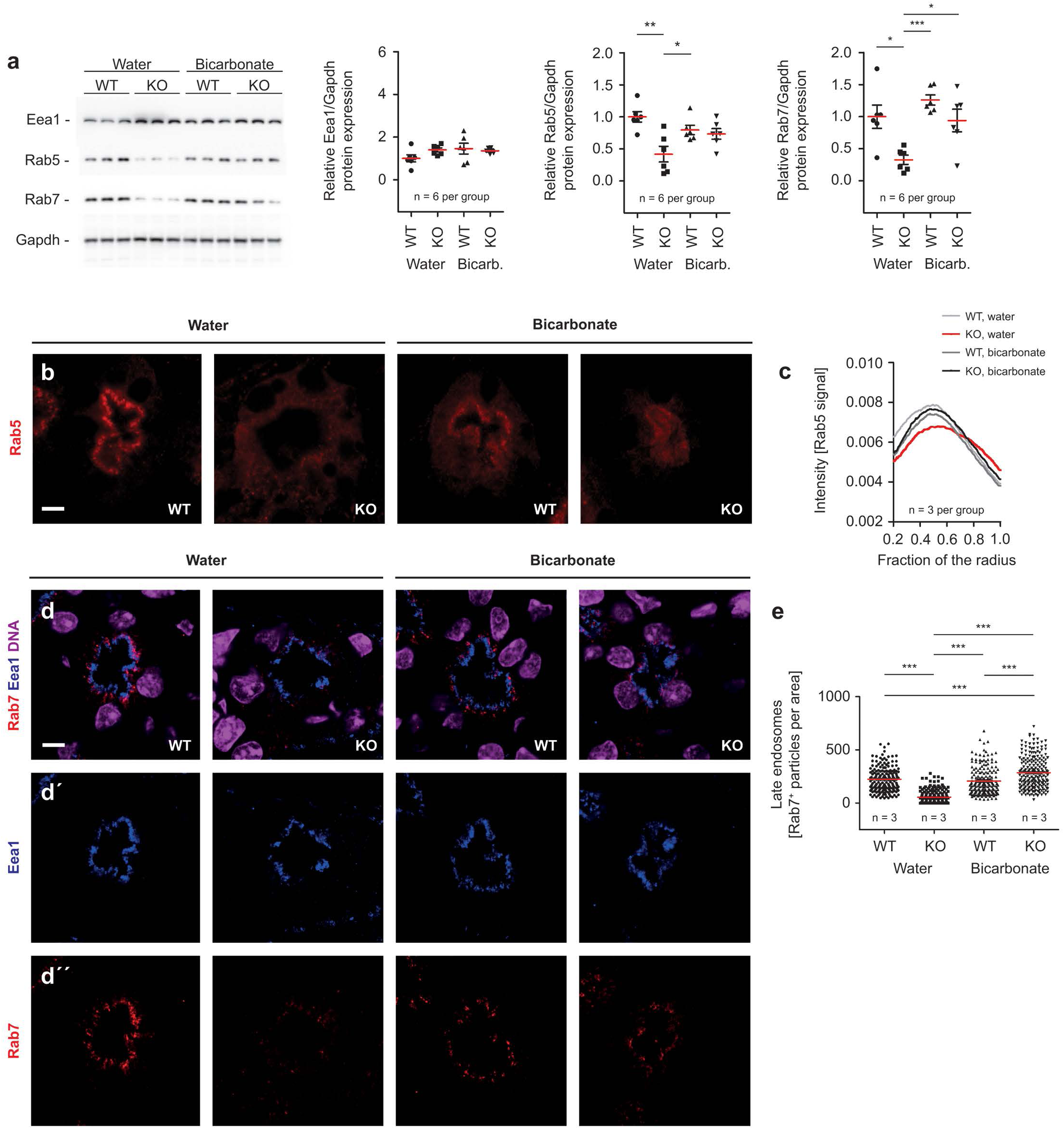
Rab7 expression is reduced in proximal tubules of *Atp6v0a4* KO mice. **(a)** The immunoblot analysis of whole kidney lysates shows no change in the abundance of Eea1, but decreased abundance of Rab5 and Rab? in *Atp6v0a4* KO mice. n = number of mice. Each data point represents one mouse. One-way ANOVA with Tukey·s multiple comparison test. **(b)** Rab5 immunofluorescence of proximal tubule sections of 21-day-old WT and *Atp6v0a4* KO mice. Scale bar 5 µm. (c) Radial intensity distribution analysis for Rab5 reveals that signal intensities are flattened in proximal tubules of untreated 21-day-old *Atp6v0a4* KO mice. n = number of mice. Mean values are shown. **(d)** Merge of Eea1 and Rab? immunofluorescence of proximal tubule sections of 21-day-old WT and *Atp6v0a4* KO mice. Scale bar 5 µm. Nuclei are stained with TO-PRO-3. **(d’)** Eea1 single channel. **(d”)** Rab? single channel. **(e)** The quantification shows a severe decrease of Rab7-labeled puncta in proximal tubules of *Atp6v0a4* KO mice. N = number of mice. Each data point represents one proximal tubule. One-way ANOVA with Tukey·s multiple comparison test. Quantitative data are represented as mean± s.e.m. with individual data points. *\*P* < 0.05; *\*\*P* < 0.005; *\*\*\*P* < 0.0005.

In summary, we show that components of the endocytic pathway are severely altered under metabolic acidosis with a depletion of Rab7-labeled later intermediates.

### Defective lysosomes and accumulation of autophagic vesicles in proximal tubules in untreated *Atp6v0a4* KO mice

Because Rab7 is a key regulatory protein for the biogenesis and maintenance of lysosomes ^23^, we studied the lysosomal compartment. Suggesting a lysosomal abnormality, the abundance of both the lysosomal protein Lamp1 and Ist1, one of the ESCRT subunits previously used to mark damaged lysosomes ^24^, were increased in untreated *Atp6v0a4* KO mice, but normal in bicarbonate-supplemented *Atp6v0a4* KO mice (**Figure 4a**). When we quantified puncta labeled for Ist1 and Lamp1 in proximal tubules, these were strongly increased in untreated *Atp6v0a4* KO mice compared to all other cohorts (**Figure 4b-c, Supplementary Figure S4a**). Compared to the regular subapical distribution we observed a shift of Lamp1-positive puncta towards a more central localization in untreated *Atp6v0a4* KO mice (**Figure 4d, Supplementary Figure S5a-c**) ^25^.

**Figure 4:**
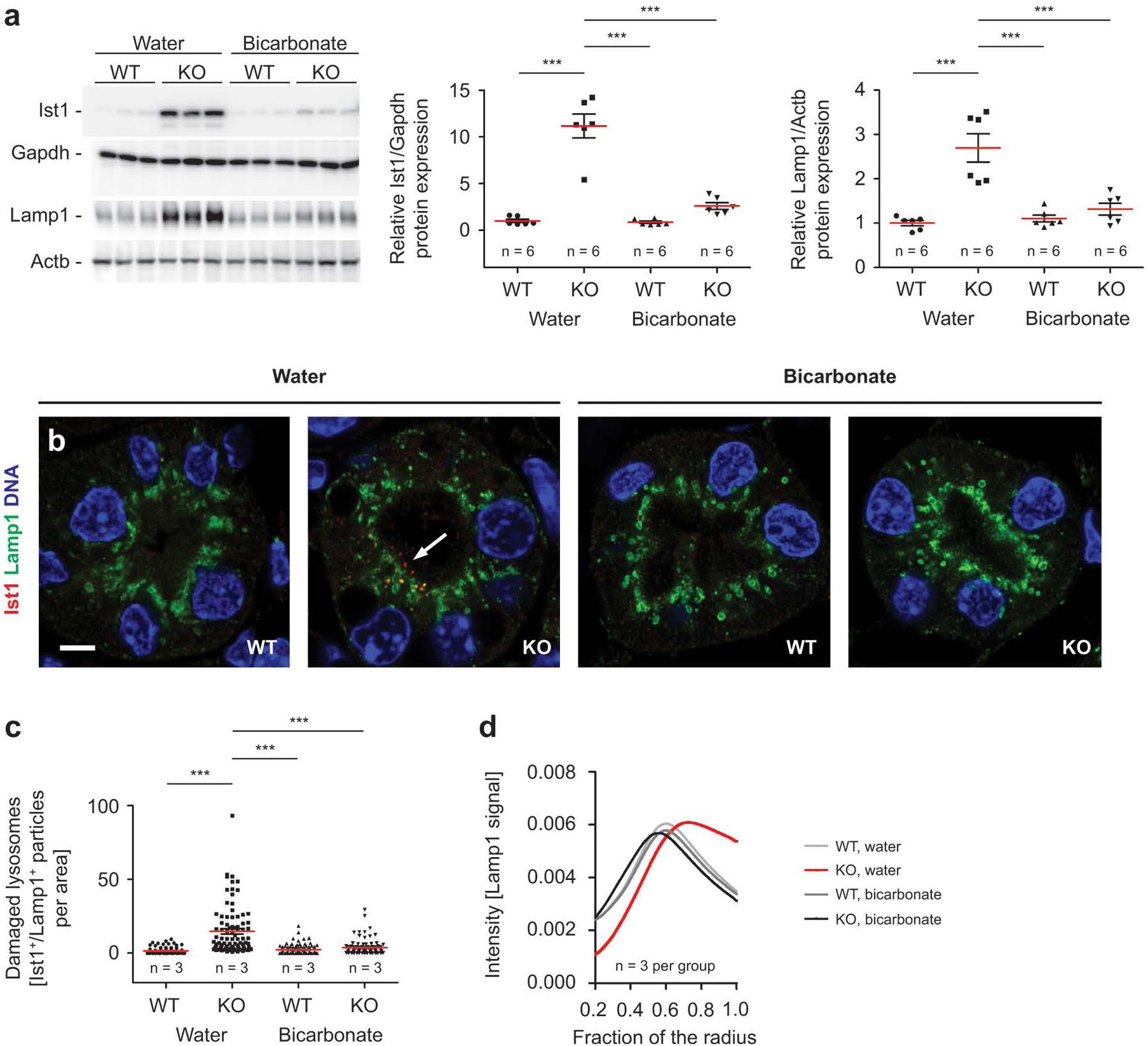
Metabolic acidosis increases the number of lst1-labeled lysosomes and alters the positioning of lysosomes in proximal tubule cells. **(a)** lmmunoblot analysis of whole kidney lysates shows an increased abundance for lst1 and Lamp1 normalized to Gapdh or Actb in untreated 21-day-old *Atp6v0a4* KO mice. n = number of mice. Each data point represents one mouse. One-way ANOVA with Tukey·s multiple comparison test. **(b)** Some Lamp1-positive puncta co-label for lst1 in proximal tubules (arrow) of 21-day-old untreated *Atp6v0a4* KO mice. No co-labeling was observed in bicarbonate supplemented *Atp6v0a4* KO mice. Single channels are shown in supplemental figures. Scale bar 5 µm. **(c)** Quantification of lst1/Lamp1-double positive puncta. n = number of mice. Each data point represents one proximal tubule. One-way ANOVA with Tukey’s multiple comparison test. **(d)** Radial intensity distribution analysis for Lamp1-positive puncta reveals that lysosomal positioning is altered in proximal tubules of untreated 21-day-old *Atp6v0a4* KO mice. n = number of mice. Mean values are shown. Quantitative data are represented as mean± s.e.m. with individual data points. *\*P* < 0.05; *\*\*P* < 0.005; *\*\*\*P* < 0.0005.

The lysosomal waste disposal system digests endocytosed material as well as cellular material through autophagy. The abundance of lipidated Map1lc3b (LC3B), which functions in autophagy substrate selection and autophagosome biogenesis, was increased in kidney lysates of untreated *Atp6v0a4* KO mice (**Figure 5a**). In accordance, we observed an accumulation of autophagosomes and autolysosomes in untreated *Atp6v0a4* KO mice compared with WT or *Atp6v0a4* KO mice supplemented with bicarbonate (**Figure 5b-e**).

**Figure 5:**
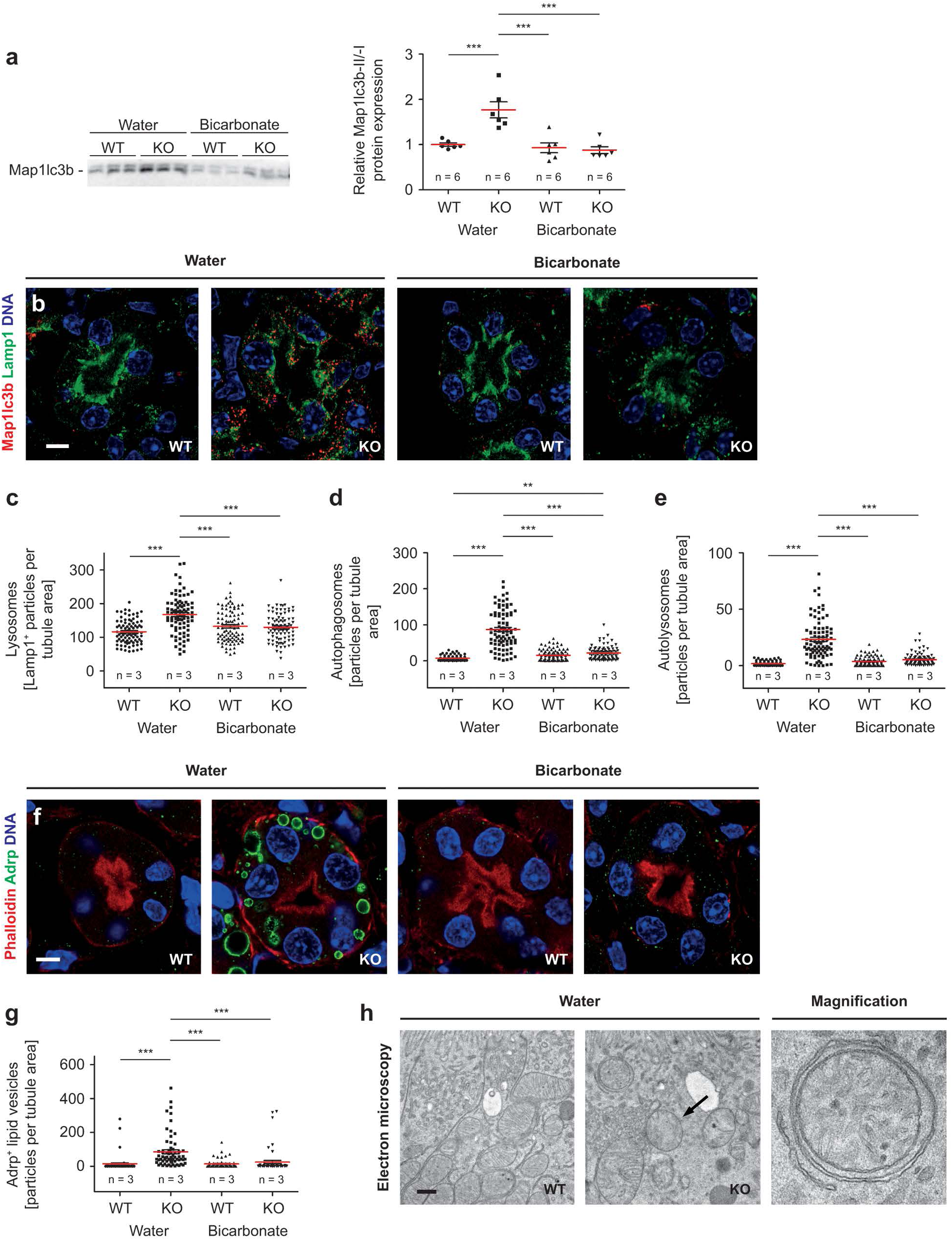
Accumulation of autophagic substrates and intermediates in proximal tubules of *Atp6v0a4* KO mice. **(a)** lmmunoblot analysis of whole kidney lysates shows increased abundance of lipidated Map1lc3b in 21-day-old untreated *Atp6v0a4* KO mice. Bicarbonate supplementation normalizes Map1lc3b abundance in *Atp6v0a4* KO mice. n = number of mice. Each data point represents one mouse. One-way ANOVA with Tukey·s multiple comparison test. **(b)** Autophagosomes (Map1lc3-positive/Lamp1-negative puncta) and autolysosomes (Map1lc3b-/Lamp1-double positive puncta) accumulate in proximal tubules of 21-day-old untreated *Atp6v0a4* KO mice and are largely absent upon bicarbonate supplementation. Scale bar 5 µm. (c) Quantification of lysosomes (Lamp1-positive puncta). n = number of mice. Each data point represents one proximal tubule. One-way ANOVA with Tukey·s multiple comparison test. **(d)** Quantification of autophagosomes (Map1lc3-positive, Lamp1-negative puncta). n = number of mice. Each data point represents one proximal tubule. One-way ANOVA with Tukey·s multiple comparison test. **(e)** Quantification of autolysosomes (Map1Ic3b- and Lamp1-double positive puncta). n = number of mice. Each data point resembles one proximal tubule. One-way ANOVA with Tukey·s multiple comparison test. **(f)** Adrp-positive structures accumulate in proximal tubules of untreated *Atp6v0a4* KO mice and are absent upon bicarbonate supplementation. Scale bar 5 µm. **(g)** Quantification of Adrp-positive lipid vesicles. n = number of mice. Each data point represents one proximal tubule. One-way ANOVA with Tukey·s multiple comparison test. **(h)** Damaged/swollen mitochondria (arrow) and mitophagy intermediates (magnification) in proximal tubules of untreated 21-day-old *Atp6v0a4* KO mice. Scale bar 500 nm. Quantitative data are represented as mean± s.e.m. with individual data points. *\*P* < 0.05; *\*\*P* < 0.005; ***P < 0.0005.

Because lipid droplets are typical substrates of autophagy ^26^, we stained for Adipophilin (Adrp), a marker of lipid droplets. Suggesting a general defect of autophagy, lipid droplets were massively increased in convoluted segments and to a lesser extent in straight segments of proximal tubules of untreated *Atp6v0a4* KO mice compared to WT or bicarbonate supplemented KO mice (**Figure 5f-g, Supplementary Figure S6a-d**). Moreover, we noticed damaged mitochondria and mitophagy intermediates in EM images of proximal tubule cells pointing towards defective mitochondria turnover in untreated *Atp6v0a4* KO mice (**Figure 5h, Supplementary Figure S7a**).

Thus, proximal tubules of acidotic *Atp6v0a4* KO mice show severe alterations of the degradative pathway with an increase in abnormal lysosomes and accumulation of autophagy substrates and intermediates.

### Disruption of the *Atp6v0a4* subunit does not perturb proximal tubule function in a cell-intrinsic manner

V-ATPase activity is essential for vesicular trafficking along the endocytic pathway of eukaryotic cells ^27, 28^. To assess whether the disruption of its a4 subunit results in a cell-intrinsic defect in proximal tubules, we generated conditional *Atp6v0a4* mice (**Supplementary Figure S8**) ^7^ utilizing Cre-recombinase under control of the rat PEPCK promoter (Hprt1^tm1(Pck1–cre)Vhh)11^. Proximal tubule-specific *Atp6v0a4* KO mice had a strictly normal acid-base status as evidenced from blood pH and bicarbonate (**Figure 6a-b**) and showed no evidence for albuminuria (**Figure 6c**). Immunofluorescence revealed that the deletion of the Atpv0a4 subunit was only mosaic (between 10 and 50%), which allowed us to directly compare proximal tubule cells with or without expression of the Atp6v0a4 subunit within the same tubule. Importantly, Lamp1 signals and their localization did not differ between deleted and undeleted proximal tubule cells (**Figure 6d**).

**Figure 6:**
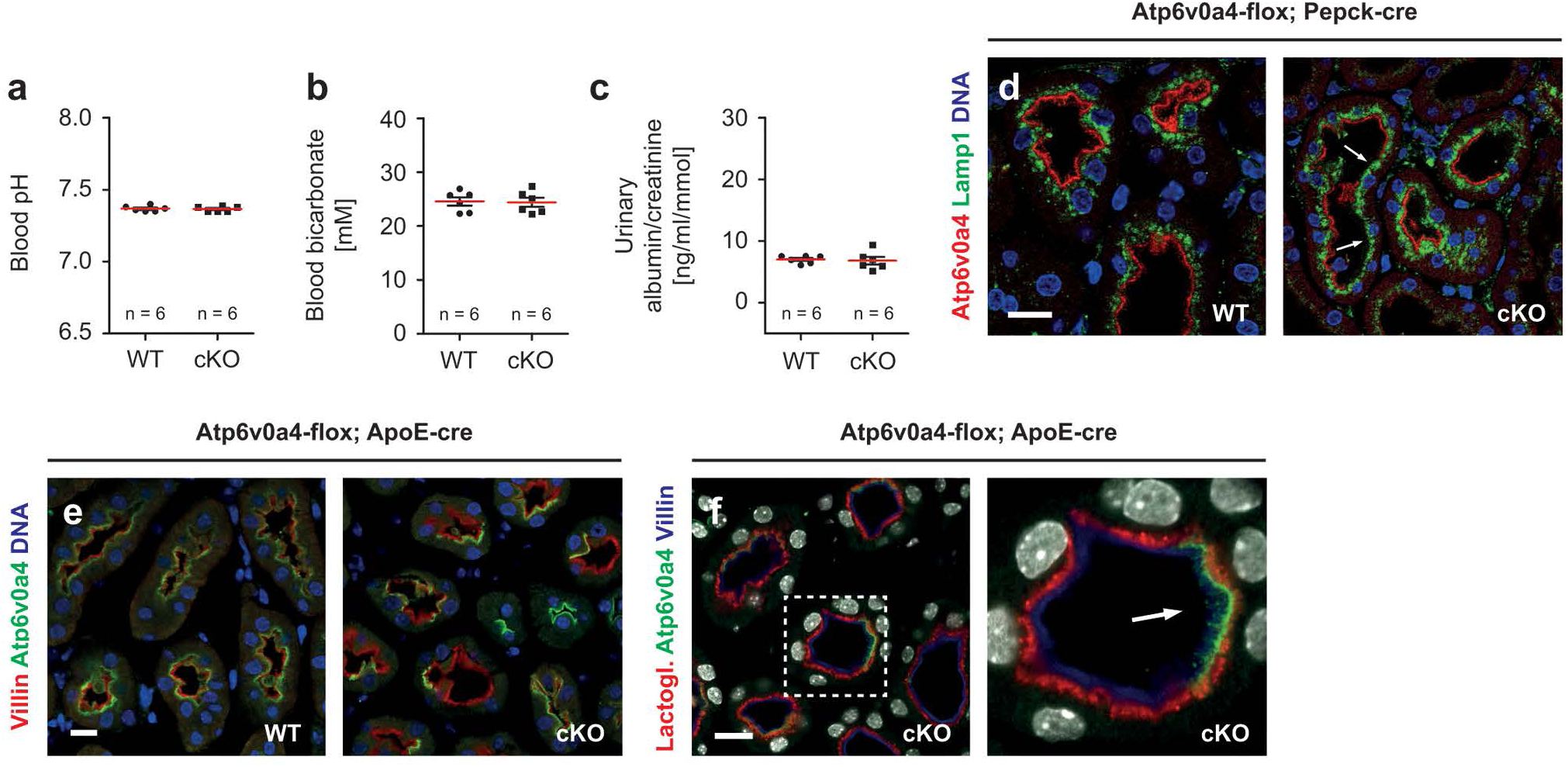
No alterations of the lysosomal compartment in proximal tubule cells of conditional cell-specific *Atp6v0a4* KO mice. (a-b) Compared to control mice (Pepck-cre·; *Atp6v0a4’^0^’1fl^0^’),* blood pH and bicarbonate levels are not changed in Pepck-cre; *Atp6v0a4-flox* mice (Pepck-cre’; *Atp6v0a4•1flox)* at 2 months of age. n = number of mice. Each data point represents one mouse. Two-sided Student’s I-test. (c) Pepck-cre; *Atp6v0a4-flox* mice do not develop albuminuria (ELISA of urine samples collected at 2 months of age). n = number of mice. Each data point represents one mouse. Two-sided Student’s I-test. **(d)** No evidence for altered positioning of lysosomes in proximal tubule cells upon deletion of the Atp6v0a4 subunit (arrows) under control of the Pepck promoter. Scale bar 1O µm. **(e)** Mosaic deletion of the Atp6v0a4 subunit in ApoE-cre; *Atp6v0a4-flox* mice. Scale bar 1O µm. **(f)** Lactoglobulin uptake by proximal tubule cells is independent of the Atp6v0a4 subunit in 2-month-old ApoE-cre; *Atp6v0a4-flox* mice (arrow = proximal tubule cell expressing the Atp6v0a4 subunit). Scale bar 1O µm. Quantitative data are represented as mean± s.e.m. with individual data points. *\*P* < 0.05; *\*\*P* < 0.005; *\*\*\*P* < 0.0005.

Moreover, we used the ApoE-Cre mouse strain ^12^ to disrupt Atp6v0a4 in proximal tubules obtained the expected mosaic deletion of Atp6v0a4 (**Figure 6e**). To assess whether endocytosis is compromised in the absence of the Atp6v0a4 subunit and normal acid base status, we injected ApoE-KO mice with fluorescently-labelled lactoglobulin and fixed the kidneys after seven minutes. Both WT and KO proximal tubule cells accumulated substantial amounts of the protein in vesicles below the brush border of proximal tubule cells, thus excluding a major endocytosis defect in proximal tubule cells devoid of Atp6v0a4 (**Figure 6f**). Taken together, these results indicate that the proximal tubule abnormalities in *Atp6v0a4* KO mice are likely not cell-intrinsic but rather secondary to metabolic acidosis.

### Systemic acidosis impairs the endocytic-degradative pathway also in WT mice

To validate our hypothesis that proximal tubule defects observed in *Atp6v0a4* KO mice are secondary to systemic acidosis and independent of the Atp6v0a4 subunit, we challenged C57BL/6 WT mice with 0.28 M NH_4_Cl-supplemented drinking water for 28 days. Compared to control mice, acid-loaded mice developed moderate systemic acidosis (**Figure 7a**) with decreased blood bicarbonate (**Supplementary Figure S9a**), increased chloride concentration (**Supplementary Figure S9b**) and decreased urinary pH (**Figure 7b**). The drop in blood pH was much less pronounced than in *Atp6v0a4* KO mice, (ΔpH acid loading: -0.1326 and ΔpH *Atp6v0a4* KO: -0.4883). Again, systemic acidosis induced the expression of key molecules required for ammoniagenesis (**Supplementary Figure S9c**).

**Figure 7:**
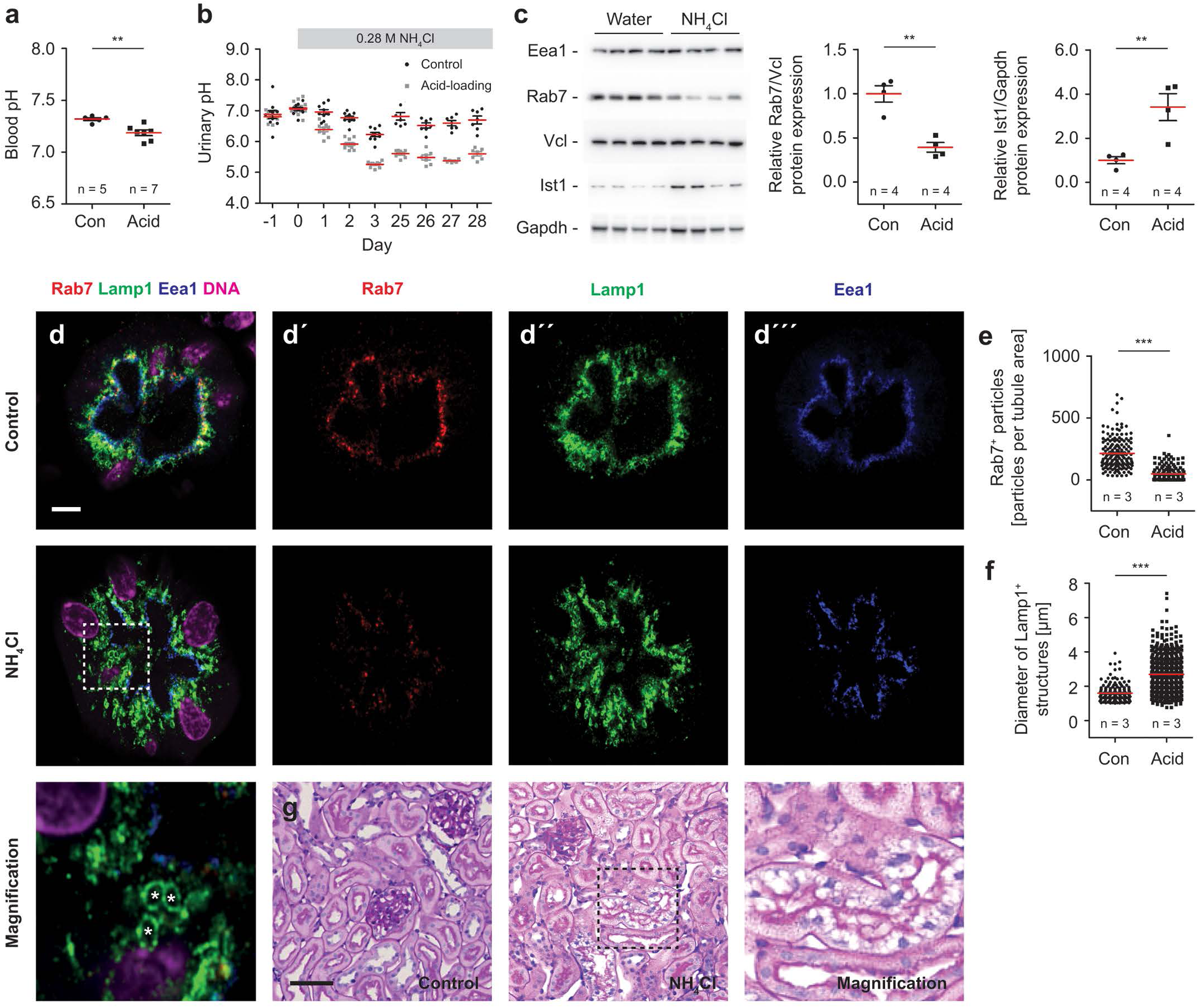
Proximal tubule trafficking defects upon acid-loading of C57BL/6 WT mice. **(a)** Addition of ammonium chloride to the drinking water for 28 days decreases blood pH in C57BL/6 WT mice. n = number of mice. Each data point represents one mouse. Two-sided Student’s I-test. **(b)** Urinary pH decreases in C57BL/6 WT mice during acid-loading. Each data point represents one mouse. (c) lmmunoblot analysis of whole kidney lysates shows decreased abundance of Rab? in acid-challenged WT mice, while lst1 abundance is increased and Eea1 abundance remains unchanged. n = number of mice. Each data point represents one mouse. Two-sided Student’s I-test. **(d)** Merge of Rab?, Lamp1 and Eea1 immunofluorescence signals in proximal tubule sections of control and acid-loaded C57BL/6 WT mice. Scale bar 5 µm. **(d’)** Rab? single channel. **(d”)** Lamp1 single channel. **(d’”)** Eea1 single channel. **(e)** Decreased number of Rab7-positive puncta in proximal tubules of acid-loaded WT mice. n = number of mice. Each data point represents one proximal tubule. Two-sided Student’s I-test. **(f)** Increased diameter of Lamp1-positive structures in proximal tubules of acid-loaded WT mice. n = number of mice. Each data point represents one proximal tubule. Two-sided Student’s I-test. **(g)** H&E staining with PAS counterstaining of kidney sections after 60 days of acid-loading compared to controls. Prolonged acid-loading results in a vacuolar degeneration of a subset of proximal tubules. Scale bar 50 µm. Quantitative data are represented as mean± s.e.m. with individual data points. *\*P* < 0.05; *\*\*P* < 0.005; ***P < 0.0005.

As in untreated *Atp6v0a4* KO mice, we observed no change in the abundance of Eea1 but a severe reduction in the abundance of Rab7 in kidney lysates from acid-loaded WT mice, while Ist1 abundance was increased (**Figure 7c**). In accordance, the number of Rab7-positive puncta was decreased in proximal tubule cells (**Figure 7d-e**). The changes in the lysosomal compartment were less pronounced compared to *Atp6v0a4* KO mice. While we observed an increase in the size of Lamp1-positive structures (**Figure 7d and f**), the distribution of lysosomes as determined by radial intensity analysis did not differ between cohorts (**Supplementary Figure S9d**). Importantly, however, we found an increased number of puncta labeled for Ist1 and Lamp1 in proximal tubules of acid-loaded mice (**Supplementary Figure S10a-b**).

After 30 days of acid loading, only very few proximal tubules showed vacuolar degeneration (**Supplementary Figure S11a-b**). At day 60, the fraction of degenerating cells in convoluted tubule segments had strongly increased (**Figure 7g and Supplementary Figure S11c-d**).

These data show that metabolic acidosis compromises proximal tubule function in general.

### Human kidney organoids recapitulate acidosis-induced defects of the endocytic pathway in proximal tubule-like structures

Because immortalized HK-2 cell lines did not fully recapitulate our *in vivo* findings and primary cultures of acutely dissociated proximal tubule cells rapidly dedifferentiated upon acid challenge (own data not shown and ^29^), we cultured kidney organoids either at control (pH 7.4), or acidic (pH 6.2) conditions for up to 3 days (**Figure 8a-d**). Within one day, we noted a downregulation of RAB7 (**Figure 8e-f and Supplementary Figure S12a-b**), an increase of the mean lysosome diameter and lysosomal damage as judged from the quantification of IST1 and LAMP1 co-labelled vesicular structures (**Figure 8g-h and Supplementary Figure S13a**) in acid challenged proximal tubule-like structures. Already two days later, we noted a drastic loss of proximal tubule-like structures in organoids cultured in acidic medium and vacuolar degeneration (**Figure 8i and Supplementary Figure S14a**).

**Figure 8:**
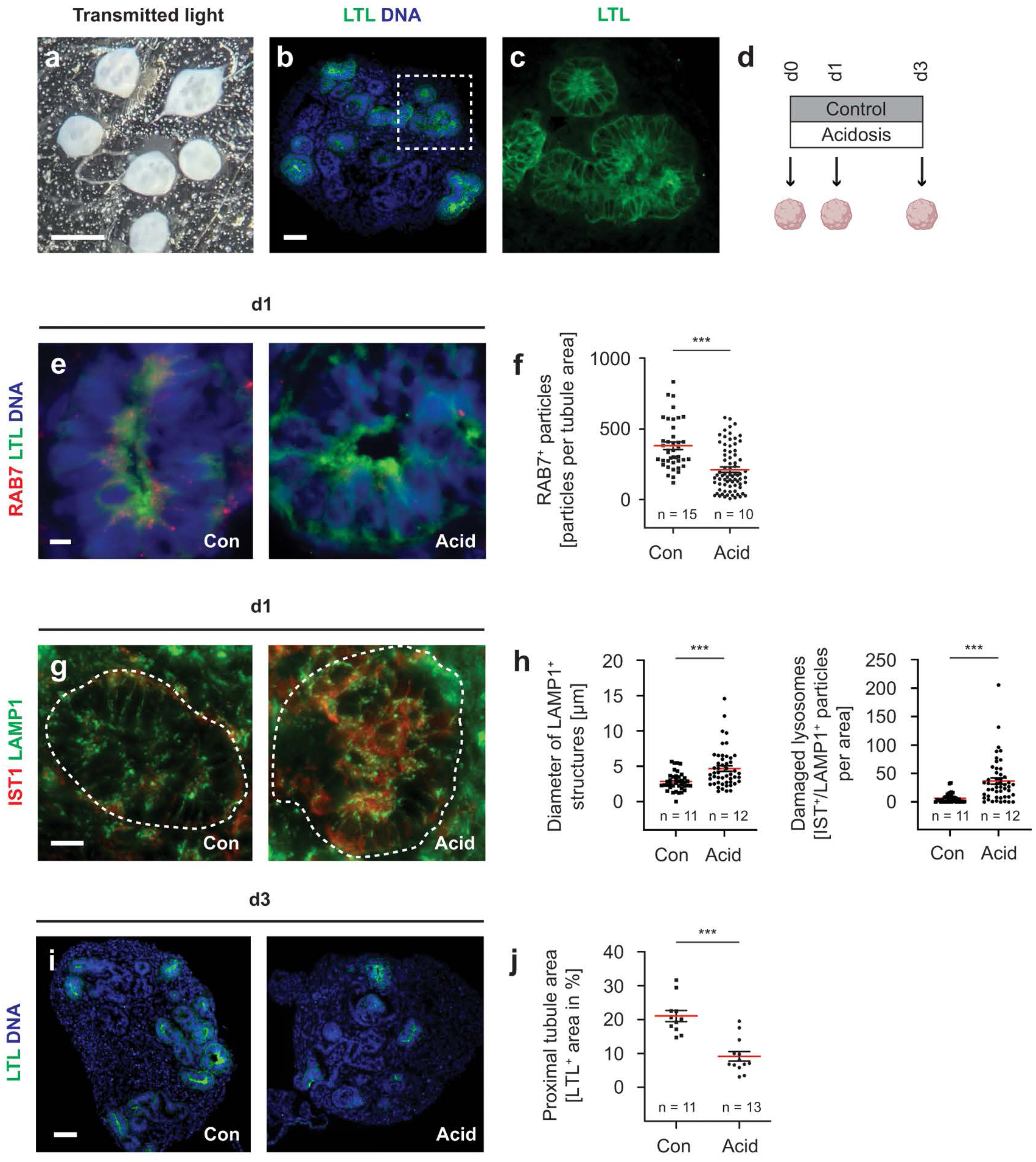
Proximal tubule-like structures in human kidney organoids recapitulate defects in the endocytic pathway during acidosis. **(a)** Transmitted light image of human kidney organoids (white structures) grown as single organoids in 96-well plates and pooled before cryo-embedding. Scale bar 500 µm. **(b)** Proximal tubule cells labelled with FITC-conjugated Lotus tetragonolobus lectin (LTL) in a human kidney organoid. Scale bar 50 µm. (c) Magnification of LTL-positive proximal tubule cells. **(d)** Schematic representation of the experiment: kidney organoids were either cultured with control (pH 7.4) or acidotic (pH 6.2) medium and fixed, stained and analyzed at the timepoints indicated. **(e)** The expression of RAB? is reduced in proximal tubule cells after 1 day of acidosis. Scale bar 5 µm. **(f)** Quantification of particles positive for RAB? per tubule area reveals reduced RABY-positive particles after 1 day of acidosis. n = number of organoids. Each data point represents one proximal tubule. Two-sided Student’s I-test. **(g)** Numbers of IST1/LAMP1-double positive particles in proximal tubules (outlined with dashed line) are increased after 1 day of acidosis. Scale bar 10 µm. **(h)** Increased diameter of LAMP1-positive structures and increased IST1/Lamp1-double positive puncta in proximal tubules after 1 day of acidosis. n = number of organoids. Each data point represents one proximal tubule. Two-sided Student’s I-test. (i) Human kidney organoids show reduced numbers of LTL-positive proximal tubules after 3 days of acidosis. Scale bar 50 µm. (j) Quantification of the proximal tubule area reveals reduced numbers of LTL-positive cells after 3 days of acidosis. n = number of organoids. Each data point represents one organoid. Quantitative data are represented as mean ± s.e.m. with individual data points. Two-sided Student’s I-test. *\*P* < 0.05; *\*\*P* < 0.005; ***P < 0.0005.

We conclude that metabolic acidosis impairs proximal tubule cells in mice and humans in a similar manner.

## Discussion

We show that severe acidosis in *Atp6v0a4* KO mice results in a Fanconi-like syndrome as evidenced by proteinuria and phosphaturia, a finding also reported for some patients suffering from dRTA ^3^. Proximal tubule dysfunction in *Atp6v0a4* KO mice is characterized by the accumulation of lipid droplets, albumin-positive deposits and autophagic vesicles and, like in human patients, is alleviated by bicarbonate supplementation. This suggests that systemic acidosis has a major impact on the degradative pathway of proximal tubule cells. Notably, the degradative pathway appeared intact in two independent mouse models with mosaic proximal tubule-specific disruption of the Atp6v0a4 subunit without systemic acidosis.

Under normal conditions, proximal tubule cells metabolize fatty acids by β-oxidation ^30–32^. Upon metabolic acidosis, however, proximal tubule cells switch to catabolism of glutamine to increase ammonia and ultimately renal acid excretion. Thus, decreased β-oxidation of fatty acids may lead to the observed increase in the number of lipid droplets in proximal tubules of untreated *Atp6v0a4* KO mice ^33^. We considered that the accumulation of lipids may also point to a defect of autophagy, because lipid droplets are a common substrate for autophagy ^34^. Supporting our hypothesis that metabolic acidosis disturbs autophagy, we observed increased lipidation of Map1lcb3 in kidney lysates and increased numbers of autophagosomes and autolysosomes in proximal tubule sections *Atp6v0a4* KO mice. Autophagy may be compromised because of the drastic increase in the number of damaged lysosomes as indicated by labeling for Ist1, one of the ESCRT subunits ^35^. In accordance with a previous report of a transient downregulation of Rab7 in the proteomic profiling of the apical membrane of the proximal convoluted tubule during onset of acidosis ^36^, we also found a strong down-regulation of Rab7 in acidotic *Atp6v0a4* KO mice. This small GTPase is fundamental for lysosomal biogenesis, the positioning and function of lysosomes, and endocytic trafficking ^37^ and central for the normal progression of the autophagic pathway in mammalian cells ^38^. The down regulation may also interfere with the recycling of proteins to the brush border such as Slc34a1. Rab7 depletion led to mis-sorting of lysosomal hydrolases and lysosomal dysfunction^39^, because it is involved in the recycling of the cation-independent mannose-6-phosphate-receptor, which delivers newly synthesized lysosomal hydrolases from the *trans*-Golgi network to endosomes. Moreover, fusion events required for the maturation of autophagosomes also require Rab7 ^40^ and its knockdown blocked the fusion of autophagosomes with late endosomes and lysosomes and inhibited the physiological activation of the lysosomal enzyme cathepsin B ^41^. Notably, Rab7 also drives lipophagy by promoting the fusion of lipid droplets with autophagosomes and lysosomes ^42^ and mitophagy, as an effector downstream of Parkin ^43^. Suggesting impaired mitochondrial turnover, we identified swollen mitochondria and mitophagy intermediates in acidotic *Atp6v0a4* KO mice. Acid-challenged Atg5-deficient proximal tubular cells exhibited reduced mitochondrial respiratory chain activity, reduced mitochondrial membrane potential, and a fragmented morphology with marked mitochondrial swelling ^44^.

Remarkably, our findings resemble data obtained for the kidney specific KO of *Vps34* (*Pik3c3*) in mice. This enzyme phosphorylates phosphatidylinositol to generate phosphatidylinositol-3-phosphate, which controls a range of cellular processes including endocytosis, Rab conversion and autophagy by reversibly recruiting specific protein effectors to membranes ^45, 46^. Proximal tubule cells devoid of Vps34 also showed preserved apico-basal specification with a regular brush border but basal redistribution of Lamp1-positive late-endosomes/lysosomes, defective endocytosis, impaired lysosomal positioning and blocked autophagy ^47^.

Importantly, we observed similar alterations of intracellular trafficking with downregulation of Rab7 and more abundant Ist1-positive damaged lysosomes in acid-challenged WT mice, which ultimately led to time-dependent vacuolar degeneration of proximal tubule cells suggesting that long-term acidosis can lead to severe proximal tubule injury. To further assess whether a similar scenario applies to the human kidney, we studied the impact of acidosis on human kidney organoids, which display highly polarized proximal tubule-like structures under control conditions. Decrease of extracellular pH caused a rapid down-regulation of RAB7 and an increase in IST1-labeled damaged lysosomes in proximal tubule equivalents within one day. Already two days later, we found a significant reduction of proximal tubule-like structures compared to control conditions, further highlighting the particular sensitivity of proximal tubules to acidosis.

Taken together, we show that severe metabolic acidosis entails drastic changes in the endocytic pathway as evidenced by changes in the abundance and distribution of Rab5 and Rab7. Because the removal of Rab5 and its replacement with Rab7 is an essential step in the formation of late endosomes and in the transport of cargo to lysosomes ^19^, metabolic acidosis may compromise the clearance of endocytosed proteins and lipids as well as autophagy, which may ultimately lead to energy deprivation despite induced glutamine metabolism. Energy deprivation will also affect the transmembrane sodium gradient generated by the basolateral Na^+^/K^+^-ATPase and thus compromise all transport processes fueled by the sodium-gradient. Thus, our observations provide insights into the mechanism by which correction of metabolic acidosis can prevent proximal tubule dysfunction. Despite in part controversial reports ^48^, it will be important to assess, whether CKD progression can be delayed by early alkali administration, as metabolic acidosis is a known risk factor for CKD progression ^49^. Moreover, it will be interesting to study, whether bicarbonate treatment has a protective or curative effect on drug-induced Fanconi syndrome.

## Disclosure

The authors declare no competing interests.

## Acknowledgements

We thank Katja Dörfel, Nicole Krönke, Johanna Fischer and Katrin Schorr for excellent technical assistance. We are indebted to Eugene Katrukha for generous help with the radial intensity distribution analysis. We thank Jörg Biber (Institute of Physiology, University of Zürich, Switzerland) for providing the Slc34a1 antibody. C.A.H. was supported by DFG grants (HU 800/7-2 and HU 800/13-1 and -2 (FOG 2625)). R.C. was supported by grants from the French National Research Agency (ANR-16-CE14-0031-01) and from the French endowment fund Philancia. D.E. was supported by a grant from the French National Research Agency (ANR-14-CE12-0013-01) and by the National Center for Precision Diabetic Medicine – PreciDIAB. (ANR-18-IBHU-0001). N.P. was supported by a grant from the French National Research Agency (ANR-16-CE140031-01[PROSTARGET], ANR-15-CE14-0024[CONTARKID], and ANR-21-CE14-0040[FATEFORA]). K.S.M. was supported by a DAAD scholarship. M.E.K. was supported by a scholarship from the IZKF Jena. B.S. was supported by the DFG (SCHE 1562/8-1 and 1562/11-1) and by the German Federal Ministry of Research and Education (BMBF grant 01GM1903B; Neocyst consortium).

## Data Sharing

All data generated or analyzed in this study were included in the main document or supplementary information files of this article.

## Supplementary Material

### Supplementary Methods

#### Randomization and blinding

For all animal experiments, mice were randomly assigned to cohorts. For the bicarbonate supplementation experiment, pregnant heterozygous mice, which had been covered by a heterozygous male mouse, were arbitrarily assigned to the control or supplementation group. This assignment was maintained for the resulting offspring. For the acid challenge in WT mice, pairs of male mice with identical body weight were split for the control and the NH_4_Cl-challenged cohort. Experimenters were not blinded for the genotype/treatment during mouse handling. For immunofluorescence studies the analyzing experimenter was blinded. Therefore, pictures were acquired and labeled in a genotype/treatment independent fashion by one experimenter and analyzed by another experimenter (KSM and JCH, respectively).

#### Renal function

Blood samples from adult mice were collected by retro-orbital bleeding under isoflurane anesthesia. Blood samples from Atp6v0a4 KO mice at 3 weeks of age were collected from anesthetized mice prior to the transcardial perfusion. All blood samples were analyzed with the ABL80 FLEX BASIC blood-gas analyzer (Radiometer). Urine samples were collected in metabolic cages (body weight > 15 g) or by voiding (body weight < 15 g). Mice destined for metabolic cages were housed in single cages. Urine pH was measured with a pH microelectrode (InLab Ultra-micro pH, Mettler-Toledo) ^1^. Urinary phosphate and creatinine were measured by HPLC ^2^ . Urinary albumin was measured with an ELISA (ICL).

#### RNA isolation, cDNA synthesis and quantitative RT-PCR

Snap-frozen kidney tissue was homogenized with mortar and pestle in liquid nitrogen. RNA was extracted with TRIzol reagent (Invitrogen) and subsequently purified with the Quick-RNA Miniprep Kit (Zymo). Total RNA was reversely transcribed into cDNA with the SuperScript IV Reverse Transcriptase (Invitrogen), random primer (Invitrogen) and RNaseOUT Inhibitor (Invitrogen). Quantitative RT-PCR was performed with commercially available TaqMan real time gene expression assays (Thermo): Slc38a3 (Mm01230670), Glutaminase (Mm01257297), Glud1 (Mm00492353), Actb (Mm00607939) on a CFX Duet Real-Time PCR System (BIO-RAD). Relative changes in gene expression were calculated with the ΔΔCt-Method with consideration of PCR efficiency ^3^.

#### Protein isolation and Western Blot

For protein isolation, snap-frozen kidney tissue was homogenized with mortar and pestle in liquid nitrogen. The kidney tissue was then homogenized on ice in 1x RIPA buffer (10 mM NaF, 150 mM NaCl, 1% NP-40, 0.5% Na-deoxycholate, 0.1% SDS, 25 mM Tris-HCl pH 7.5, 1 mM EDTA pH 8.0) with cOmplete proteinase inhibitor (Roche) and PhosSTOP phosphatase inhibitor (Roche) using a douncer. Insoluble debris was centrifuged in a cooled centrifuge and the protein lysate completed by addition of Laemmli buffer and denaturation at 60°C for 10 minutes. Protein lysates were separated with Tricine-SDS-PAGE and blotted onto PVDF membranes ^4^. Membranes were blocked with 5% non-fat dry milk (Santa Cruz) in 1x TBST for one hour at room temperature (RT), rinsed in 1x TBST and incubated with primary antibodies in 0-5% Bovine Serum Albumin Fraction V (Serva) in 1x TBST overnight at 4°C. Membranes were washed with 1x TBST, incubated with HRP-coupled secondary antibodies for 2 hours at RT and washed with 1x TBST before development with Clarity Western ECL substrate (BIO-RAD) in an LAS4000 (Cytiva). The quantification of band intensities was performed with ImageJ/Fiji (NIH).

#### Histology

Toluidine blue and Periodic acid-Schiff (PAS) staining were described in ^5^. HE staining followed standard protocols as reported in ^6^.

#### Tissue fixation and processing for electron microscopy

Kidneys were perfusion-fixed via the abdominal aorta. 3% hydroxyethyl starch in 0.1 M sodium-cacodylate (Caco buffer) was injected for 20 to 30 seconds followed by 3% paraformaldehyde, 3% hydroxyethyl starch in Caco buffer for 5 minutes. Tissues were post-fixed overnight at RT in 1.5% glutaraldehyde, 1.5% paraformaldehyde, 0.05% picric acid in Caco buffer, then in 1% osmium tetroxide, 0.8% potassium hexacyanoferrate in Caco buffer for 1.5 h at RT for TEM or in 1% aqueous osmium tetroxide for SEM, then dehydrated and embedded in epoxy resin for semithin sectioning and light microscopy (LM) or ultrathin sectioning and TEM analysis using standard methodology.

#### Electron microscopy

1 µm semithin sections were prepared from Epon blocks with an ultramicrotome (Ultracut E, Reichert-Jung) and stained with Richardson’s stain for LM evaluation. Ultrastructural analyses were performed using a Zeiss EM901 TEM equipped with a digital camera system.

#### Tissue fixation and immunofluorescence

In general, all immunofluorescence experiments were performed on sections of at least three independent mice per cohort. If not indicated otherwise, representative images are displayed in the figures. Kidneys were fixed by transcardial perfusion with 4% paraformaldehyde in 1x phosphate buffered saline (1x PBS) of with Xylazine (16 mg/kg body weight) and Ketamine (100 mg/kg body weight) deeply anesthetized mice. Fixed kidney tissue was cryoprotected with 30% sucrose in 1x PBS, embedded in Tissue-Tek O.C.T. (Sakura) and frozen on dry ice. 12 µm tissue sections were cut with a Cryotome (Thermo). Before immunostaining, tissue sections were transferred into Shandon cassettes (Sequenza) and antigen retrieval was performed in 1% SDS, 1x PBS for 5 minutes. Tissue sections were blocked with 5% NGS, 1x PBS for 1 hour at RT. Primary antibody incubation was performed overnight at 4°C. After washing the sections with 1x PBS, secondary antibodies were incubated for 2 hours at RT. Before mounting with Fluoromount-G mounting medium (Invitrogen), tissue sections were washed and either stained with DAPI (Invitrogen) or TO-PRO-3 stain (Thermo), depending on the previously utilized fluorophore-coupled secondary antibodies. Stained tissue sections were analyzed with a Zeiss LSM 880 confocal microscope.

#### Lactoglobulin injection

Mice were injected with bovine ß-Lactoglobulin (Sigma) labelled with Alexa Fluor 546 as described in ^7^.

#### iPSC culture

Human induced pluripotent stem cells (iPSCs)KOLF2.1J ^8^ were maintained in mTeSR1 media (STEMCELL Technologies, 05850) on dishes coated with Matrigel (Corning, 354277) in a 37 °C incubator with 5% CO2.

#### Differentiation of iPSCs into renal organoids

iPSCs differentiation protocol was based on Morizane et al. ^9^. Briefly, cells were dissociated into single cells using Accutase (STEMCELL Technologies, 07920) and plated into 10-cm dishes (800.000 cells/dish) coated with Matrigel (Corning, 354277) in mTeSR1 with 10 µM ROCK inhibitor Y-27632 (STEMCELL Technologies; #72304). After 36 h, cells were briefly washed with PBS and then cultured in basic differentiation medium (Advanced RPMI 1640 (Life Technologies, #12633-020) with 1x L-GlutaMAX (Life Technologies, #35050-061)) supplemented with 7 µM CHIR99021 (STEMCELL Technologies; #100 1042) for 96 h to induce late primitive streak cells. To induce posterior intermediate mesoderm, cells were then cultured in basic differentiation medium with 10 ng/ml activin A (STEMCELL Technologies; #78132.1) for 3 days. For induction of nephron progenitor cells, the media was then changed to basic differentiation medium with 10 ng/ml FGF9 (STEMCELL Technologies; #78161.1)) for 2 days.

#### 3D renal organoid formation

iPSCs on day 9 of differentiation, which represents metanephric mesenchyme cells, were dissociated with Accutase and resuspended in the basic differentiation medium supplemented with 3 μM CHIR and 10 ng/ml FGF9, and placed in 96-well, round bottom, ultra-low-attachment plates (Corning, #7007) at 5 · 10^5^ cells per well. The plates were centrifuged at 300 g for 15 s, and the cells then cultured at 37°C, 5% CO2 for 2 days. The medium was then changed to the basic differentiation medium supplemented with 10 ng/ml FGF9 and cultured for 3 more days. After that, the organoids were cultured in basic differentiation medium with no additional factors for 11 days.

#### Immunohistochemistry of renal organoids

3D kidney organoids media with indicated pH was changed every two days. Organoids were collected at day 0, 3, 5 and 7 after treatment to a 12-well plate, washed with PBS, fixed with 4% paraformaldehyde in PBS for 1h, washed three times in PBS, then incubated with 30% sucrose in PBS (w/w) overnight at 4°C. The organoids were mounted and frozen in Tissue-Tek O.C.T. compound (Sakura). 5 μm sections were washed three times in PBS, and permeabilized in 0.5% Triton X-100 in PBS, for 15 min. After blocking with 1% normal donkey serum (NDS, Biozol, END9010) and 1% bovine serum albumin (BSA, Thermo Fisher Scientific, 23 209) in PBS, primary antibody incubation was performed overnight at 4°C. The next day, sections were washed with 3x PBS and incubated with secondary antibodies for 1 hour at RT. Afterward, the organoids were mounted with Prolong gold without DAPI (Invitrogen) and subjected to immunofluorescence microscopy. Pictures were taken with a Zeiss AxioObserver microscope equipped with Colibri 7 LED light source, Axiocam 702 mono and Apotome system, C-Apochromat 63x/1.20 W objective with the Zeiss ZEN 3.3 software. Subsequent image processing and analysis were performed using ImageJ/Fiji Software version 2.9.0/1.53t (NIH).

#### Antibodies

The following antibodies were used in this study.

Actb (Proteintech, 20536-1-AP, Rabbit, 1:2500 for WB)

Adrp (Proteintech, 15294-1-AP, Rabbit, 1:500 for IF)

Alb (Proteintech, 16475-1-AP, Rabbit, 1:500 for IF)

Atp6v0a4 (Proteintech, 21570-1-AP, Rabbit, 1:500 for IF)

Eea1 (Proteintech, 68065-1-Ig, Mouse, 1:1000 for IF, 1:5000 for WB)

FcRn (Proteintech, 67944-1-Ig, Mouse, 1:1000 for IF)

Gapdh (Proteintech, 10494-1-AP, Rabbit, 1:5000 for WB)

Ist1 (Proteintech, 19842-1-AP, Rabbit, 1:200 for IF, 1:1000 for WB)

Lamp1 (BD Biosciences, #553792, Rat, 1:1000 for IF, 1:1000 for WB)

Lamp1 (Abcam, ab25630, Mouse, 1:400 for IF, organoids only)

Lrp2 (Proteintech, 19700-1-AP, Rabbit; 1:1000 for IF)

LTL (Vector laboratories, FL-1321-2, coupled with FITC, 1:500 for IF, organoids only)

Map1lc3b (NanoTools, clone 5F10, 0231-100, Mouse, 1:50 for IF)

Map1lc3b (Cell Signaling, #2775, Rabbit, 1:1000 for WB)

Rab5 (Cell Signaling, clone C8B1, #3547, Rabbit, 1:500 for IF, 1:1000 for WB)

Rab7 (Abcam, ab137029, Rabbit, 1:500 for IF, 1:5000 for WB)

Slc34a1 (NaPi-IIa, gift from Jörg Biber, Institute of Physiology, University of Zürich, University of Zürich, Switzerland)

Vcl (Cell Signaling, #4650, Rabbit, 1:1000 for WB)

Villin (Proteintech/Acris, 66096-1-lg, Mouse, 1:200 for IF)

Phalloidin (Invitrogen)

DAPI (Invitrogen)

TO-PRO3 (Invitrogen)

All fluorophore-coupled secondary antibodies (Invitrogen, 1:1000 for IF)

#### Particle and colocalization analysis with Fiji

Microscopy images from kidney sections were acquired at identical laser and microscopy settings and were equally post-processed. Using ImageJ/Fiji, the single channel pictures were transformed into 8-Bit and a threshold for signal intensity applied. The threshold function results in a pixel with either signal or no signal, which is necessary for the downstream application. The respective single channels for further analysis were merged and then subjected to the ComDet tool (Katrukha E. 2020, ComDet plugin for ImageJ, v0.5.3, Zenodo, doi:10.5281/zenodo.4281064). The ComDet tool allows the quantification of particle numbers (1 particle = 4 pixels, larger particles were segmented) and the colocalization between two markers analyzed. Roughly, a minimum of 30 random proximal tubules were analyzed per mouse.

#### Radial intensity distribution analysis

To assess the distribution of signals within proximal tubule cells from the apical towards the basolateral side, we used a radial intensity distribution analysis tool (18) of ImageJ/Fiji. The outer boundaries of entire proximal tubules were marked as circled region of interest (ROI) and the Radial intensity macro was started. Intensity projections from at least 30 tubules per mouse with 3 mice per group were analyzed.

#### Size analysis of Lamp1-positive structures in C57BL/6 mice

Proximal tubules were randomly selected and Lamp1-positive vesicular structures with a diameter larger than 1 µm were identified using ImageJ/Fiji. 3 mice per cohort were analyzed with a minimum of 10 proximal tubules per mouse.

#### Study approval

All animal experiments were approved by local authorities. <u>France:</u> Animal house agreement no. A 974 001 (Authorization for Animal Experimentation no. 201806111409218, Project approval by our local ethical committee no. CEEA 114). <u>Germany:</u> Thüringer Landesamt für Verbraucherschutz (TLV): 02-035/13, 02-038/16, 02-068/16 and UKJ-17-006. Landesamt für Gesundheit und Soziales (LaGeSo): O 0334/11.

## Supplementary Figures

**Supp. Figure S1:**
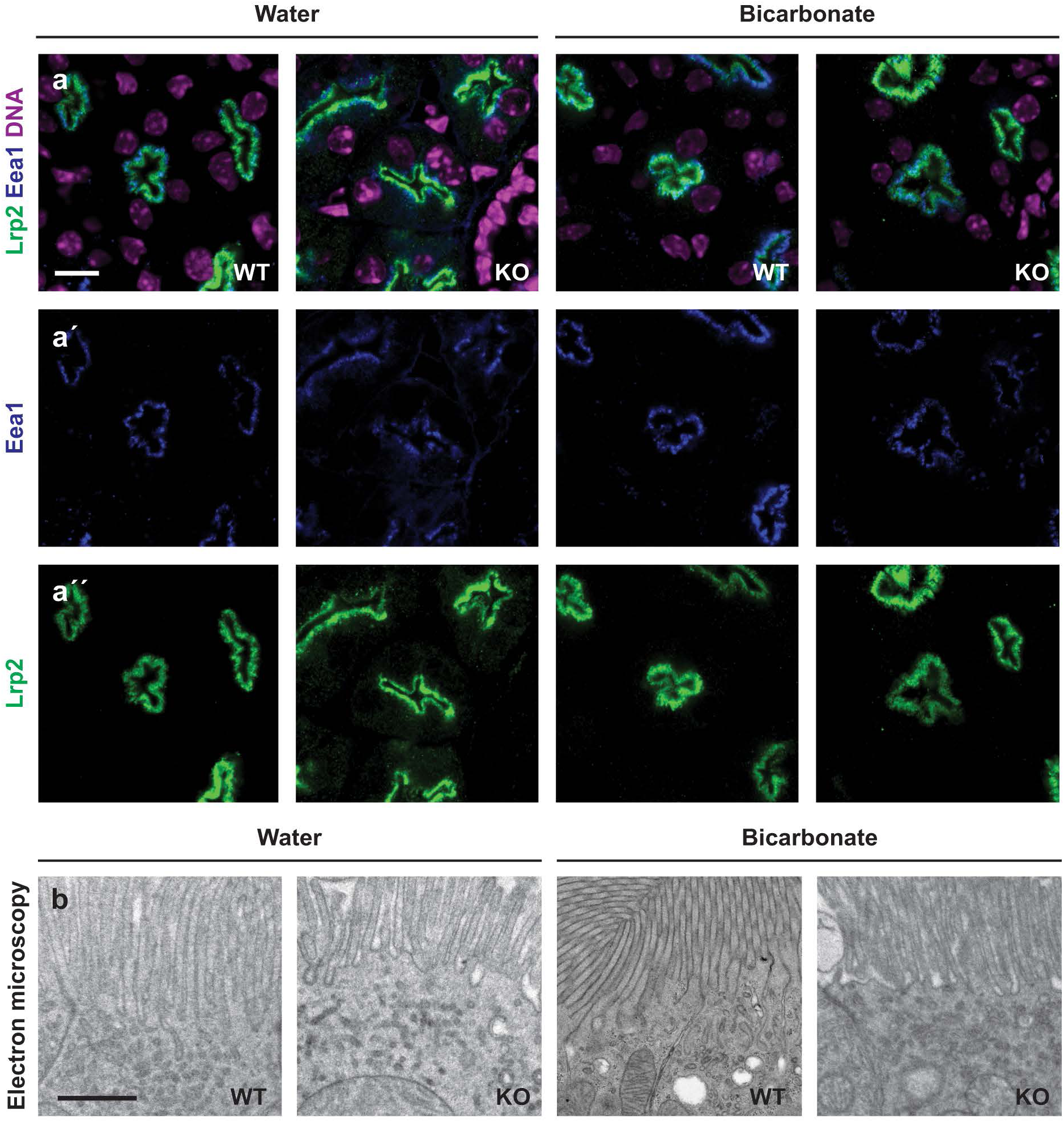
No evidence for major changes in the structure of the brush border under metabolic acidosis. **(a)** Merge of Lrp2 and Eea1 immunofluorescence signals of proximal tubule sections of 21-day-old WT and *Atp6v0a4* KO mice. Scale bar 5 µm. **(a’)** Eea1 single channel. **(a”)** Lrp2 single channel. **(b)** Electron microscopy showing the brush border membrane. Scale bar 1 µm.

**Supp. Figure S2:**
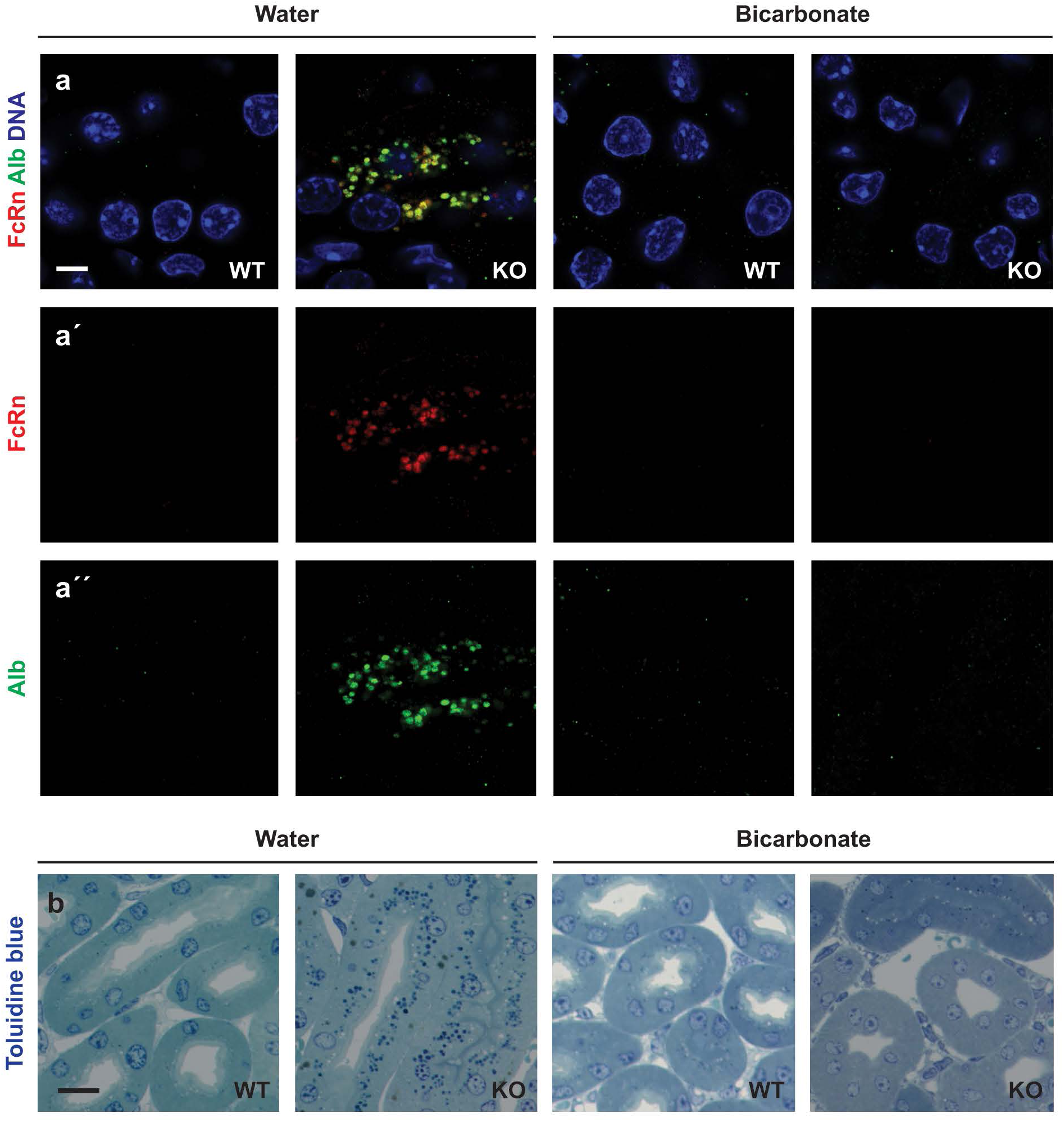
Single channels for FcRn and Alb staining. **(a)** Albumin- and FcRn-positive vesicles accumulate in proximal tubule cells in the medulla of untreated 21-day-old *Atp6v0a4* KO mice, but not in WT and/or in bicarbonate-treated mice. Scale bar 5 µm. **(a’)** FcRn single channel. **(a”)** Albumin single channel. **(b)** Toluidine blue staining shows medullary protein deposits (dark blue) in *Atp6v0a4* KO mice not supplemented with bicarbonate. Bicarbonate treatment prevents the formation of protein deposits. Scale bar 10 µm.

**Supp. Figure S3:**
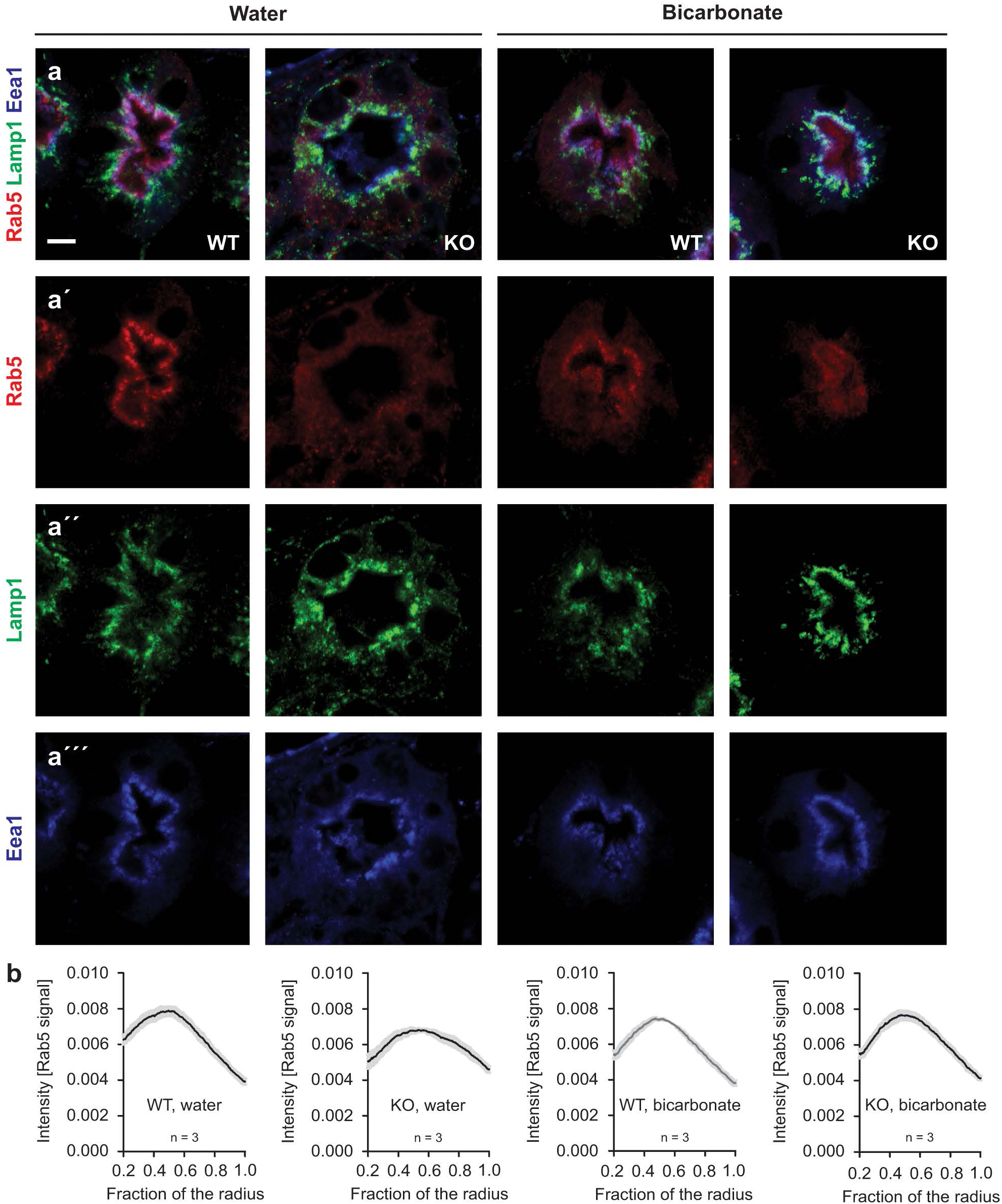
Acidosis alters the intensity profiles of Rab5. **(a)** Merge of Rab5, Lamp1 and Eea1 immunofluorescence signals of proximal tubule sections of 21-day-old WT and *Atp6v0a4* KO mice. Scale bar 5 µm. **(a’)** Rab5 single channel. **(a”)** Lamp1 single channel. **(a’”)** Eea1 single channel. **(b)** Radial intensity distribution analysis for Rab5 reveals that signal intensities are flattened in proximal tubules of untreated 21-day-old *Atp6v0a4* KO mice. n = number of mice. A minimum of 30 tubules per mouse were analyzed. Mean values with s.e.m. are shown.

**Supp. Figure S4:**
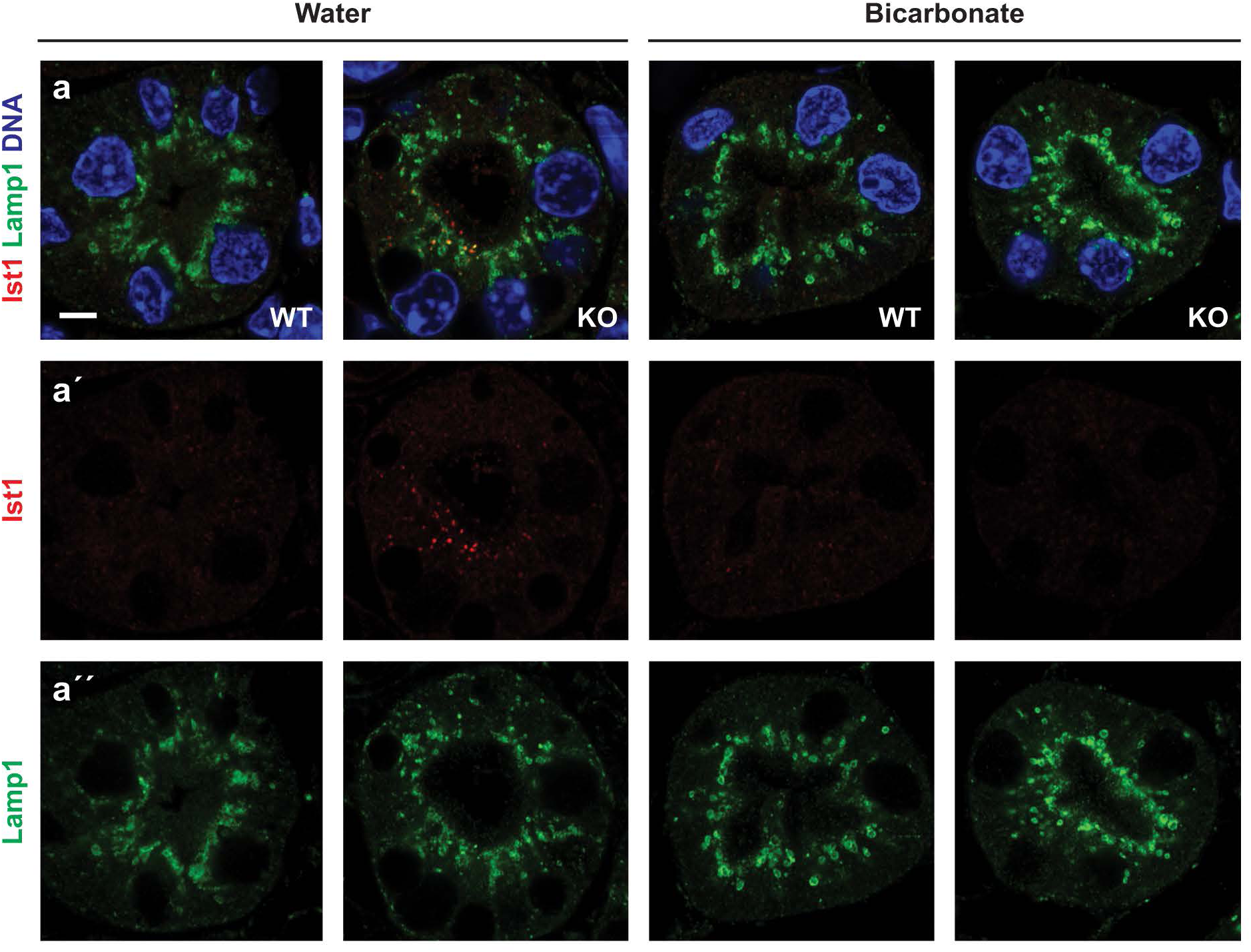
Single channels for lst1 and Lamp1 staining. **(a)** Some Lamp1-positive puncta co-label for lst1 in proximal tubules of 21-day-old untreated *Atp6v0a4* KO mice. No co-labeling was observed in bicarbonate supplemented *Atp6v0a4* KO mice. Scale bar 5 µm. **(a’)** lst1 single channel. **(a”)** Lamp1 single channel.

**Supp. Figure S5:**
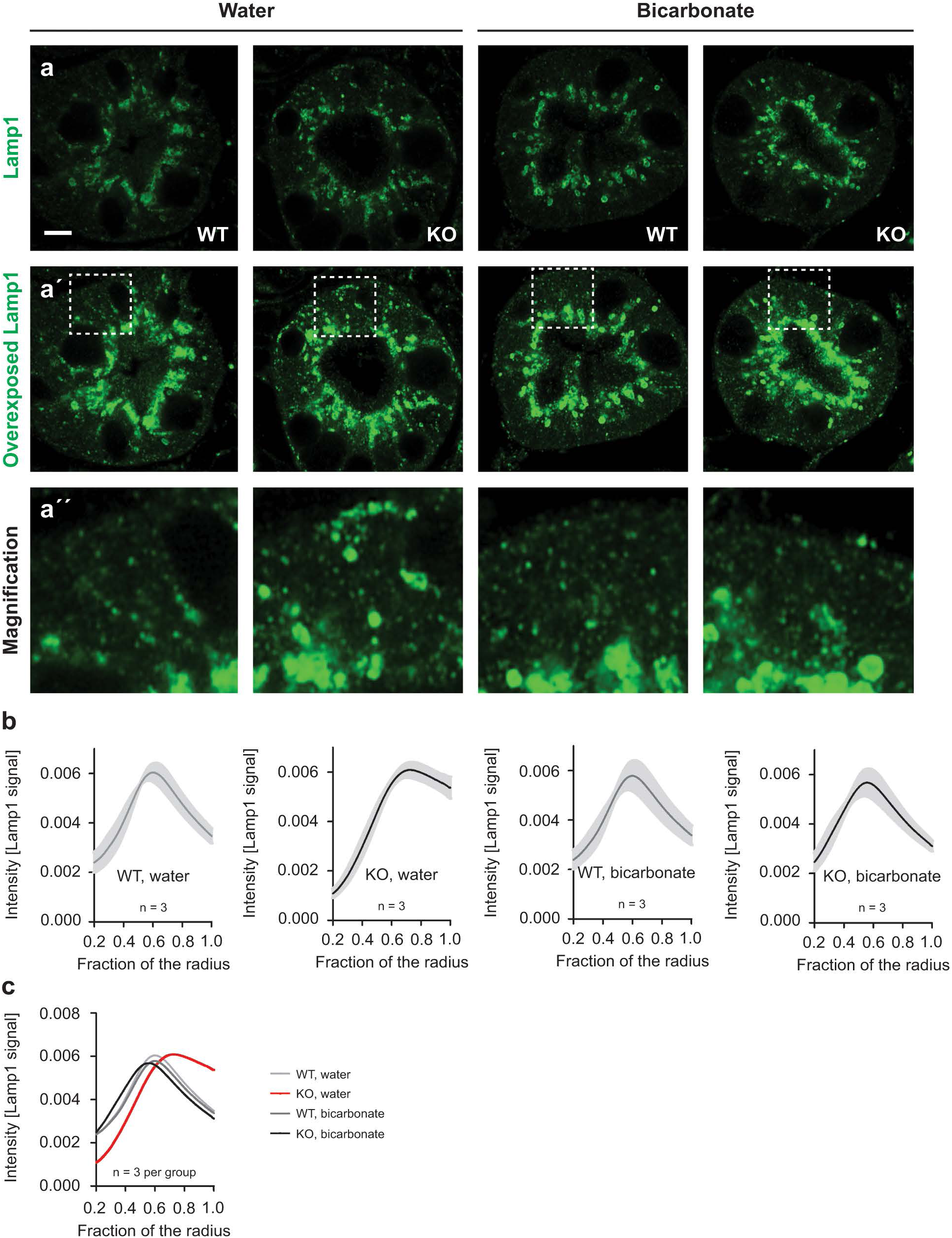
Radial intensity analysis for Lamp1. **(a)** Lamp1 single channel immunofluorescence in proximal tubules of control and *Atp6v0a4* KO mice at 21 days of age with and without bicarbonate treatment. Note: Identical tubules are shown in Figure 4 and Suppl. Figure 4. Scale bar 5 µm. **(a’)** Overexposed Lamp1 channel. **(a”)** Magnification of the overexposed Lamp1 single channel to highlight the basolateral localization of Lamp1 in untreated *Atp6v0a4* KO mice. **(b)** Radial intensity analysis for each cohort. Mean values with s.e.m. are shown. **(c)** Overlay of mean values for each cohort. n = number of mice. A minimum of 30 tubules per mouse was analyzed. Mean values with s.e.m. are shown.

**Supp. Figure S6:**
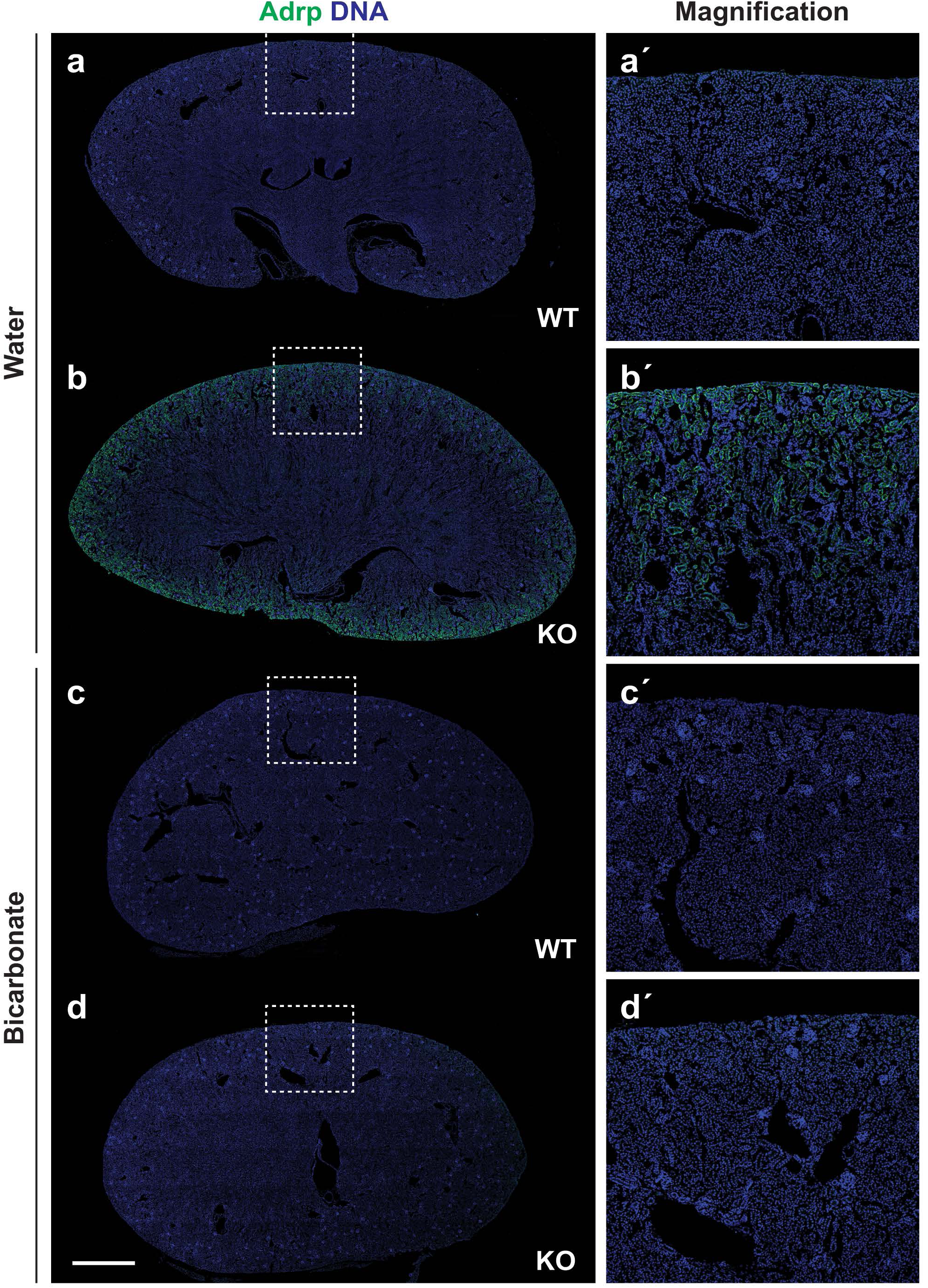
Whole kidney section overview for Adrp staining. Adrp and DAPI staining on 21-day-old whole kidney sections from untreated WT **(a),** untreated *Atp6v0a4* KO **(b),** bicarbonate-treated WT (c) and bicarbonate-treated *Atp6v0a4* KO mice **(d).** Magnification of the boxed area **(a’ -d’).** Scale bar 1 mm.

**Supp. Figure S7:**
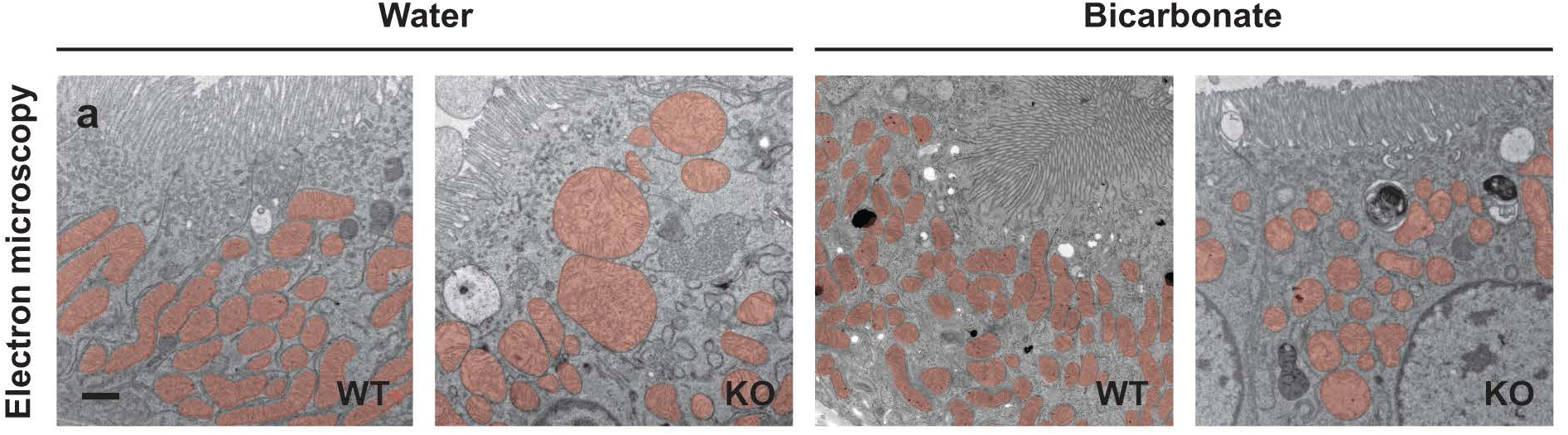
Electron microscopy of mitochondria in proximal tubules. Damaged and swollen mitochondria in proximal tubules of untreated 21-day-old *Atp6v0a4* KO mice. Mitochondria are highlighted in red. Scale bar 1 µm.

**Supp. Figure S8:**
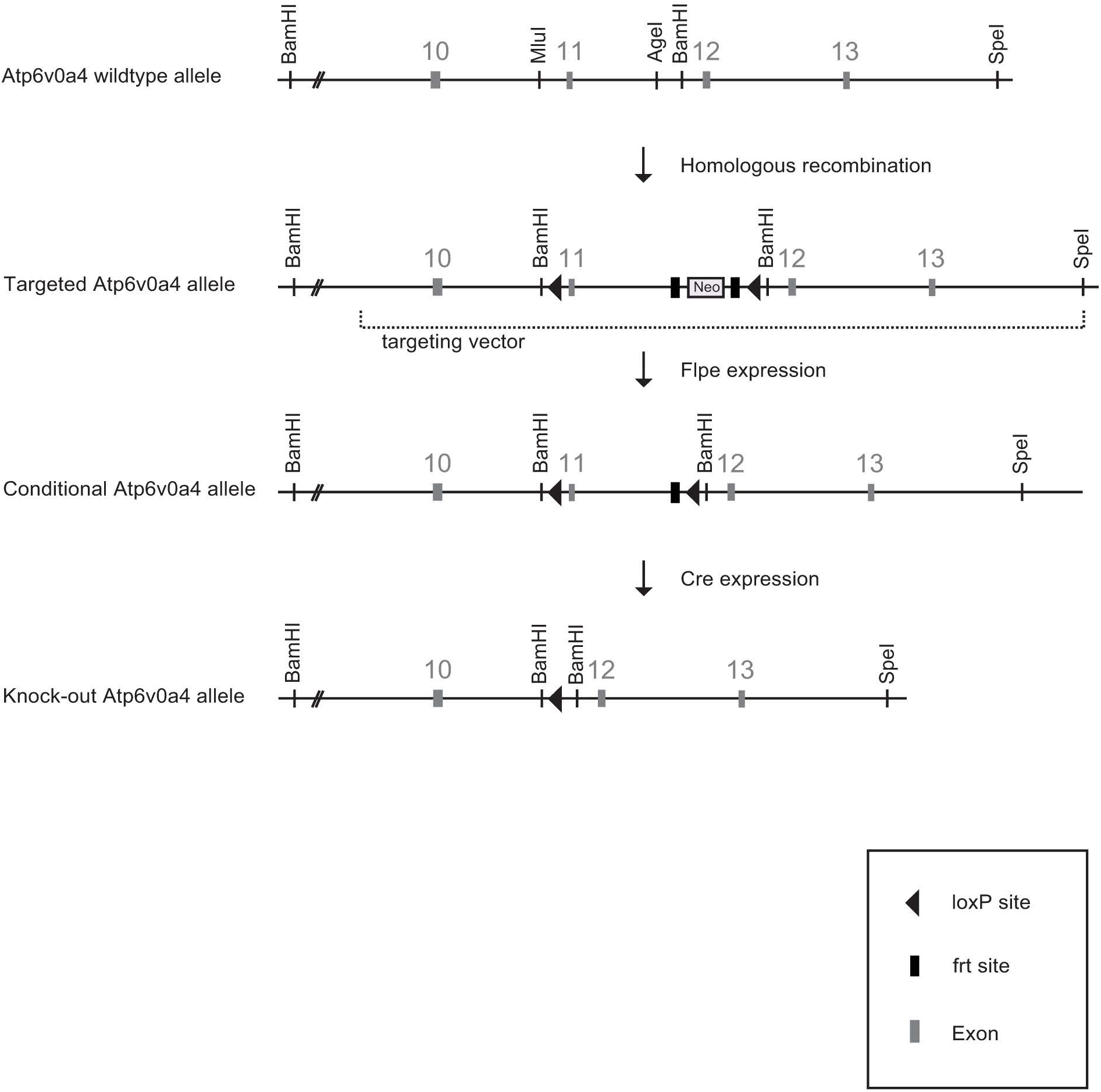
*Atp6v0a4* targeting strategy and generation of conditional *Atp6v0a4* KO mice. Homologous recombination of the targeting vector with Neomycin resistance cassette. Transient Flpe expression results in excision of the Neomycin resistance cassette. The resulting ES-cell clone was injected into blastocysts to establish the floxed line. Exon 11 is flanked by loxP sites and can be deleted by ere-expression.

**Supp. Figure S9:**
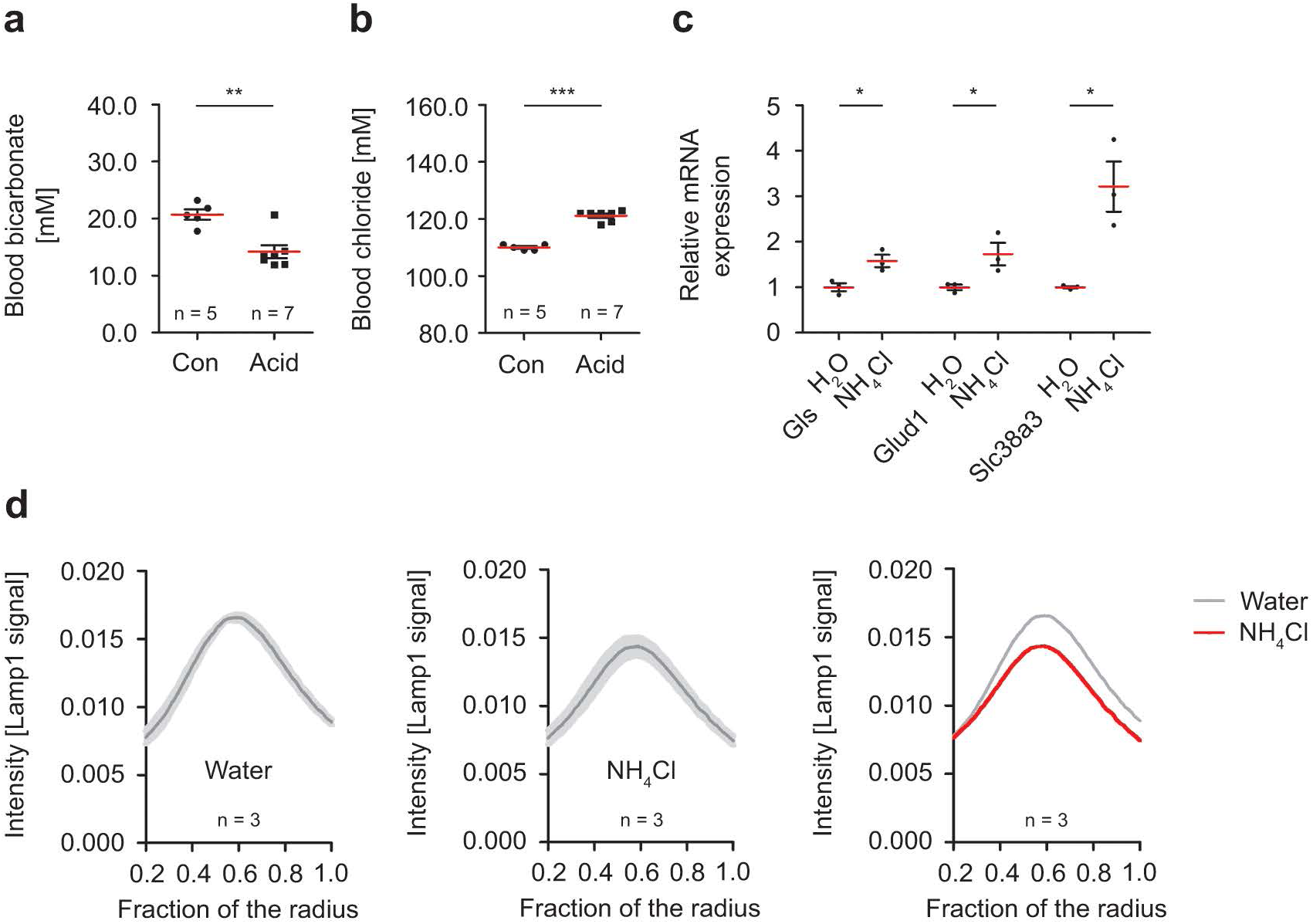
Acid-loading of C57BU6 WT mice. (a-b) Addition of ammonium chloride to the drinking water for 28 days decreases blood bicarbonate in C57BL/6 WT mice, while blood chloride is increased. Two-sided Student’s t-test. **(c)** Ammonium chloride treatment increases the expression for known ammoniagenesis genes (Glutaminase (Gls), Glutamate dehydrogenase 1 (Glud1), Solute carrier family 38 member 3 (Slc38a3)) as shown by quantitative **RT-PCR** on whole kidney tissue mRNA from 3-month-old mice. Two-sided Student’s t-test. **(d)** Radial intensity distribution analysis for Lamp1-positive puncta shows that lysosomal positioning is not altered in proximal tubules of acid-loaded C57BL/6 WT mice, but signal intensities are reduced. n = number of mice. At least 30 tubules per mouse were analyzed. Mean values are shown. Quantitative data are represented as mean ± s.e.m. with individual data points. *\*P* < 0.05; *\*\*P* < 0.005; *\*\*\*P* < 0.0005.

**Supp. Figure S10:**
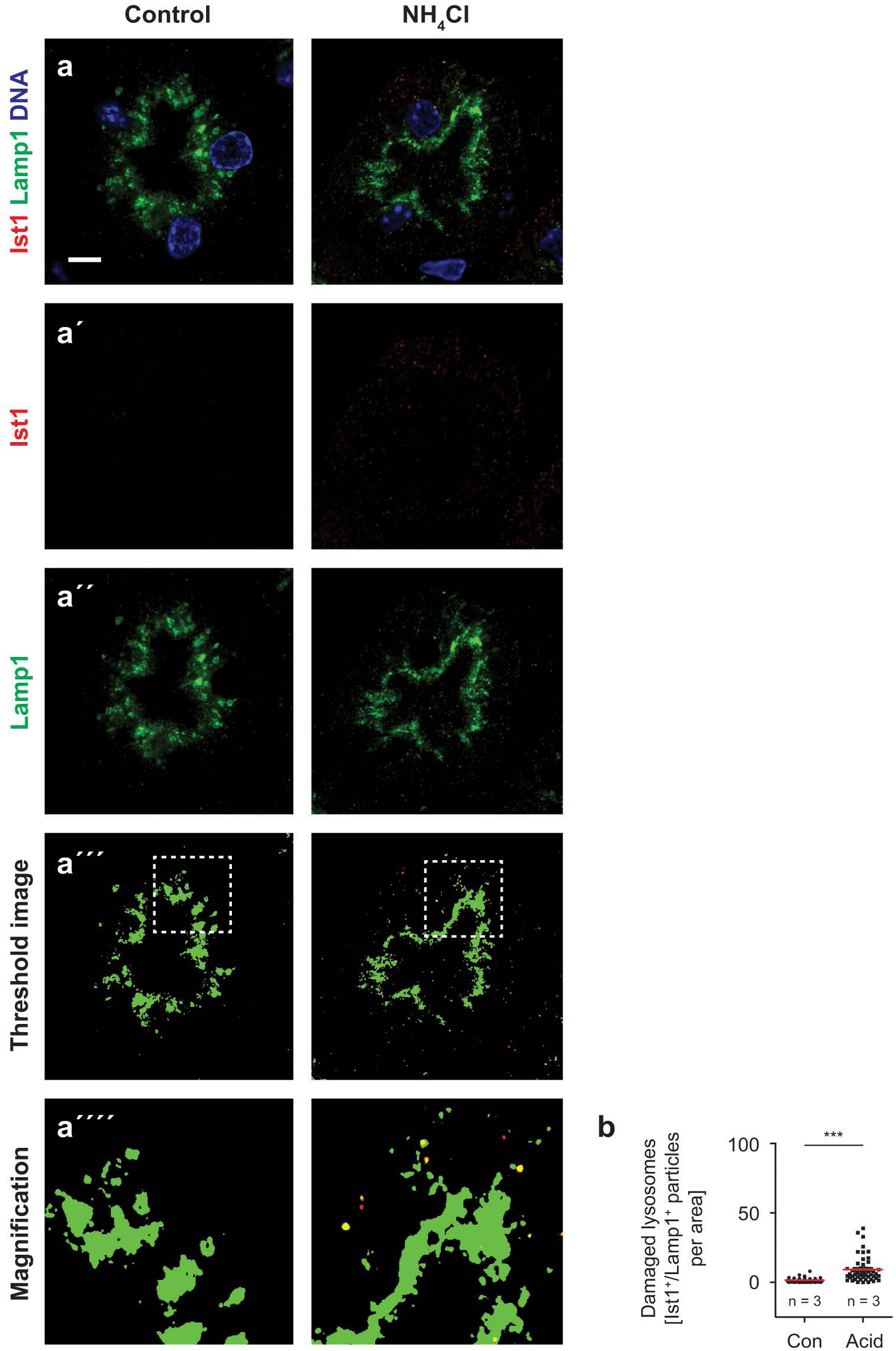
Acid-loading increases the number of damaged lysosomes in C57BU6 WT mice. **(a)** Partial colocalization of lst1 and Lamp1 in proximal tubules of acid-loaded C57BL/6 WT mice at 3 months of age. Scale bar 5 µm. **(a’)** lst1 single channel. **(a”)** Lamp1 single channel. **(a’”)** Edited picture with threshold (signal or no signal) function in lmageJ. **(a””)** Magnification of the threshold image to highlight the lst1 and Lamp1 colocalization. **(b)** Quantification of lst1/Lamp1-double positive puncta. n = number of mice. Each data point resembles one proximal tubule. Two-sided Student’s t-test. Quantitative data are represented as mean± s.e.m. with individual data points. *\*P* < 0.05; *\*\*P* < 0.005; *\*\*\*P* < 0.0005.

**Supp. Figure S11:**
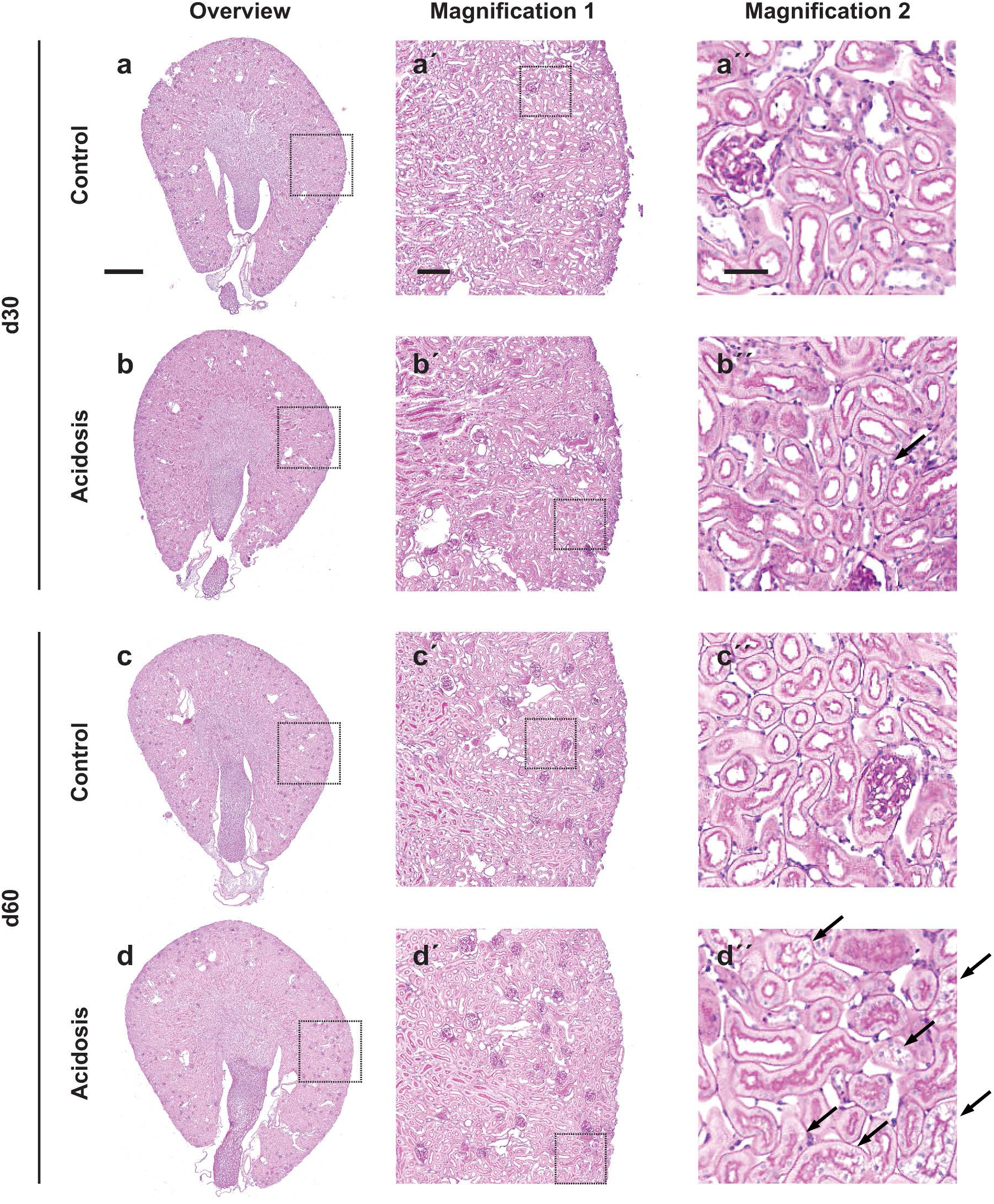
Whole kidney sections of C57BL/6 WT mice after 30 days and 60 days of acid-loading. **(a)** Whole kidney section overview of a control C57BL/6 mouse after 30 days. Scale bar 500 µm. **(a’)** Magnification of the boxed area. Scale bar 100 µm. **(a”)** Magnification of the boxed area. Scale bar 25 µm. **(b)** Whole kidney section overview of a C57BL/6 mouse after 30 days of acid-loading. A very small subset of proximal tubules shows vacuolar degeneration (arrow). **(b’)** Magnification of the boxed area. **(b”)** Magnification of the boxed area. **(c)** Whole kidney section overview of a control C57BL/6 mouse after 60 days. **(c’)** Magnification of the boxed area. **(c”)** Magnification of the boxed area. **(d)** Whole kidney section overview of a C57BL/6 mouse after 60 days of acid-loading. Many proximal tubules show vacuolar degeneration (arrows). **(d’)** Magnification of the boxed area. **(d”)** Magnification of the boxed area.

**Supp. Figure S12:**
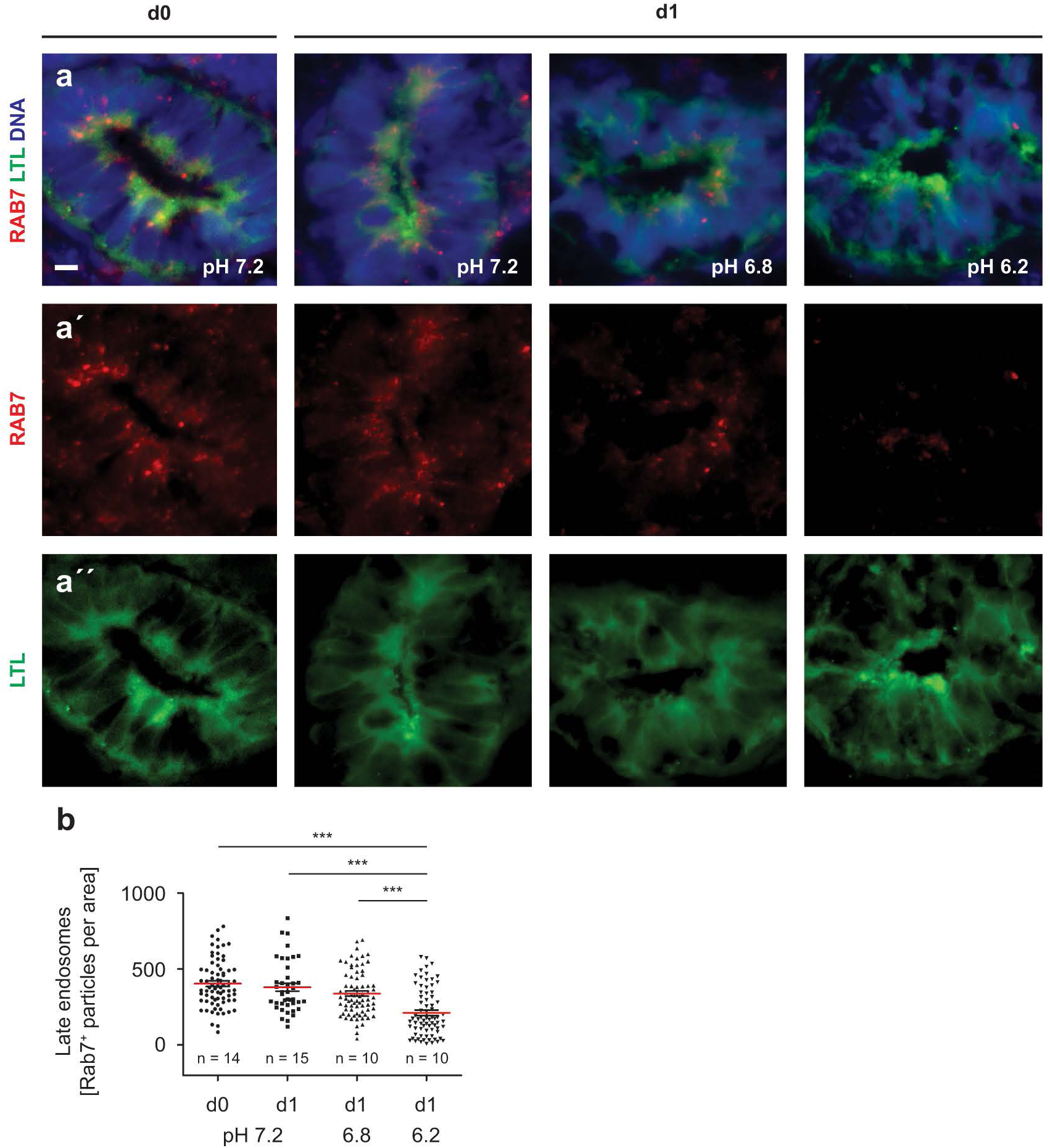
Single channels for RAB7 and LTL staining. **(a)** lmmunofluorescence staining for RAB?, **LTL** and DNA at day O and day 1. The extracellular pH was set to pH 6.8 and 6.2, respectively. Control organoids were cultured at pH 7.2. Scale bar 5 µm. **(a’)** RAB? single channel. **(a”) LTL** single channel. **(b)** The quantification of RABY-positive puncta reveals a reduction of RAB? after 1 day of acidosis. At normal pH or prior to acidification, no changes in RAB? puncta were observed. n = number of organoids. Each data point represents one proximal tubule. One-way ANOVA with Tukey’s multiple comparison test. Quantitative data are represented as mean ± s.e.m. with individual data points. *\*P* < 0.05; *\*\*P* < 0.005; *\*\*\*P* < 0.0005.

**Supp. Figure S13:**
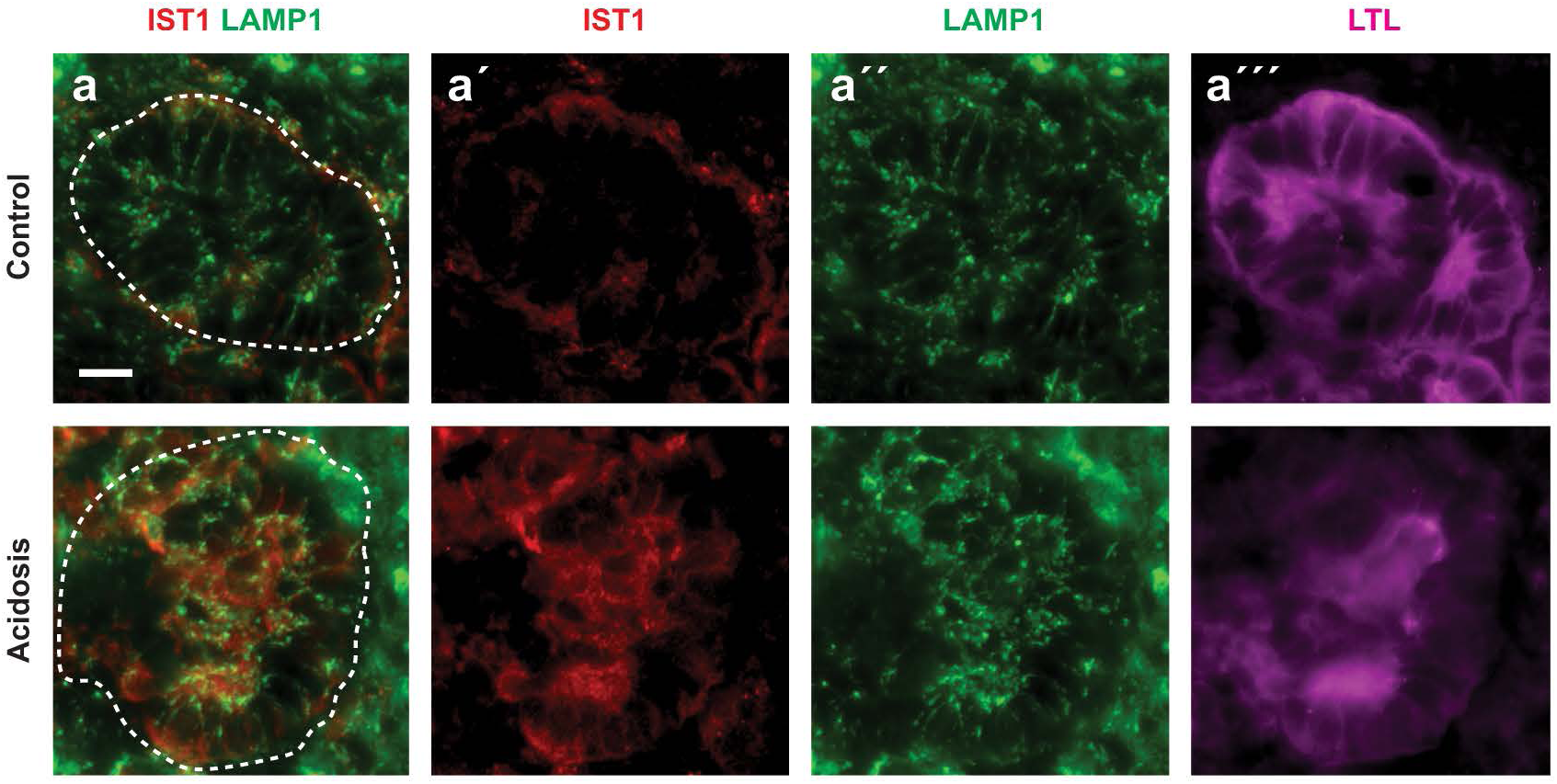
Single channels for IST1, LAMP1 and LTL staining. **(a)** Partial colocalization of IST1 and LAMP1 in proximal tubule-like structures (outlined by dashed line) of human kidney organoids at day 1. **(a’)** IST1 single channel. **(a”)** LAMP1 single channel. **(a’”)** LTL single channel. Scale bar 10 µm.

**Supp. Figure S14:**
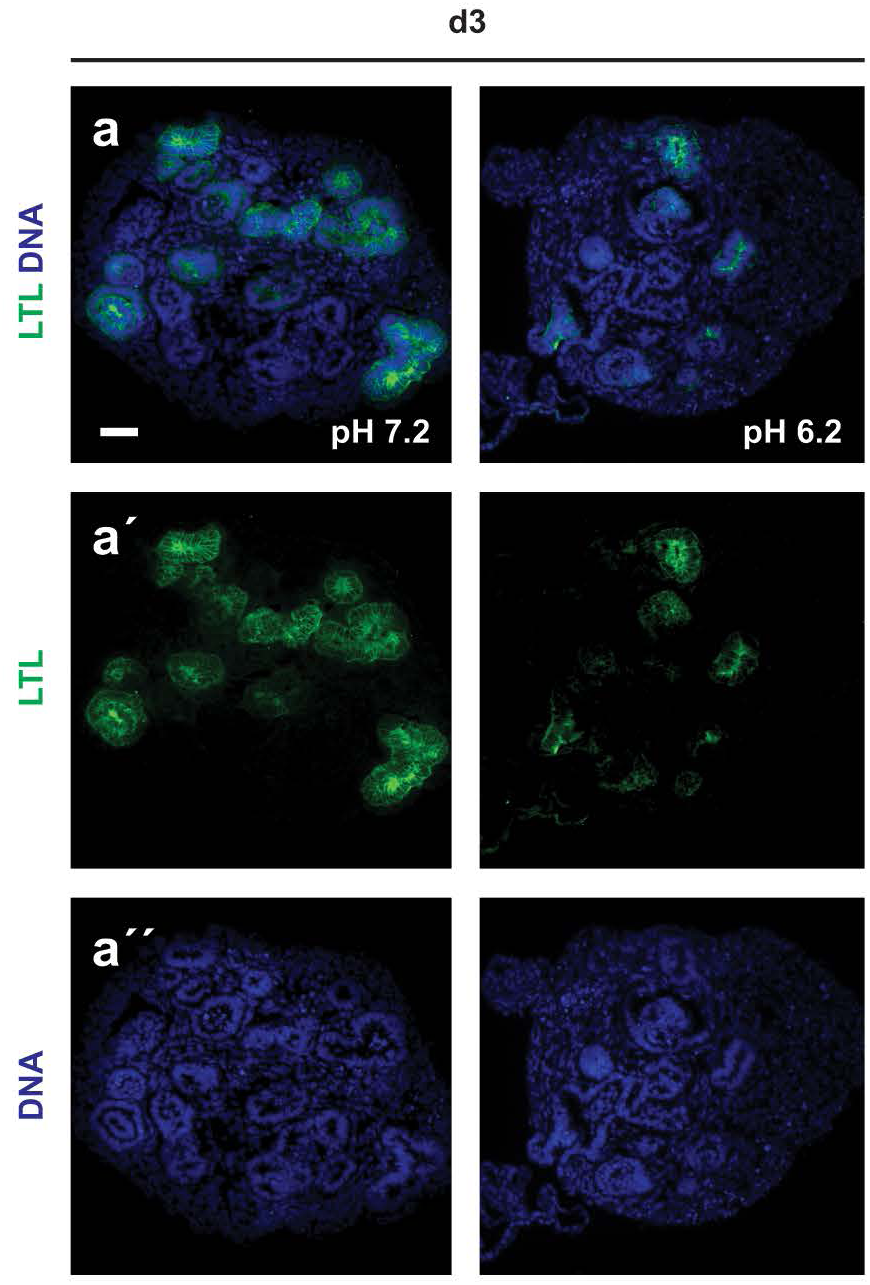
Single channels for LTL and DAPI staining. **(a)** Staining of proximal tubule-like structures with LTL in human kidney organoids at day 3. Under acidosis, proximal tubule-like structures are reduced. **(a’)** LTL single channel. **(a”)** DAPI single channel. Scale bar 50 µm.

